# A temperature-inducible protein module for control of mammalian cell fate

**DOI:** 10.1101/2024.02.19.581019

**Authors:** William Benman, Zikang Huang, Pavan Iyengar, Delaney Wilde, Thomas R. Mumford, Lukasz J. Bugaj

## Abstract

Inducible protein switches allow on-demand control of proteins in response to inputs including chemicals or light. However, these inputs either cannot be controlled with precision in space and time or cannot be applied in optically dense settings, limiting their application in tissues and organisms. Here we introduce a protein module whose active state can be reversibly toggled with a small change in temperature, a stimulus that is both penetrant and dynamic. This protein, called Melt (<u>Me</u>mbrane localization through temperature), exists as a monomer in the cytoplasm at elevated temperatures but both oligomerizes and translocates to the plasma membrane when temperature is lowered. The original Melt variant switched states between 28-32°C, and state changes could be observed within minutes of temperature change. Melt was highly modular, permitting thermal control over diverse processes including signaling, proteolysis, nuclear shuttling, cytoskeletal rearrangements, and cell death, all through straightforward end-to-end fusions. Melt was also highly tunable, giving rise to a library of variants with switch point temperatures ranging from 30-40°C. The variants with higher switch points allowed control of molecular circuits between 37°C-41°C, a well-tolerated range for mammalian cells. Finally, Melt permitted thermal control of cell death in a mouse model of human cancer, demonstrating its potential for use in animals. Thus Melt represents a versatile thermogenetic module for straightforward, non-invasive, spatiotemporally-defined control of mammalian cells with broad potential for biotechnology and biomedicine.

## Main Text

Inducible proteins permit on-demand, remote control of cell behavior, for example using chemicals or light as inputs. These inputs trigger protein conformational changes that can regulate a vast array of downstream protein and cell behaviors in a modular manner. While chemical control requires delivery of a small molecule, light can be applied remotely and offers further benefits for precision in both space and time, as well as low cost of the inducer. There is tremendous potential to extend these benefits into more complex settings including in 3D cell and tissue models, in patients for control of cell therapy, or in dense bioreactors for bioproduction. However, optical control is limited in these more opaque settings because visible light cannot penetrate, scattering within millimeters of entering human tissue^1,2^. Non-optical forms of energy like magnetic fields or sound waves can travel deeper but generally lack protein domains that can sense and respond to these stimuli. There is thus a need for protein switches that can couple penetrant and spatiotemporally precise stimuli to the control of intracellular biochemistry in living cells.

Temperature has gained recent interest as a dynamic inducer in opaque settings^3–6^. Unlike light, temperature can be readily controlled in tissues. Simple application of an ice pack or heat pad can change tissue temperature at ∼cm length scales^7^. For deeper and more precise control, focused ultrasound can be used to heat tissue with sub-millimeter-scale spatial resolution^8^. Furthermore, unlike either chemical- or light-induction, thermal-responsiveness could uniquely interface with an organism’s own stimuli, setting the stage for engineered biological systems that autonomously detect and respond to physiological temperature cues, for example fevers or inflammation.

The widespread adoption of chemo- and optogenetic proteins was enabled by protein domains that undergo stereotyped changes in response to small molecules or light. However, remarkably few analogous temperature-sensing modules have been described. Temperature-sensitive (Ts) mutants are protein variants that denature at elevated temperatures ^9–11^, but such mutants are generally neither modular nor reversible and must be laboriously validated for each individual target. The TlpA protein from *Salmonella* forms thermolabile dimers^12^ and underlies existing thermosensitive engineered proteins, including a temperature-controlled dimerization module ^13^. However, TlpA-based dimers are large (∼600-700 amino acids in combined size), and may be limited by the need for stoichiometric tuning between the two components. Elastin-like polypeptides form condensates at elevated temperatures, but these have mostly been engineered for use outside of cells, and the few intracellular applications do not have an appropriate temperature profile for use in mammalian systems^14–16^. At the level of transcription, heat shock promoters have been used for thermal control, including to induce tumor clearance by engineered cells ^4,17,18^. However heat shock promoters can respond to non-thermal stimuli ^19–21^, and thermal response profiles cannot be readily tuned because they depend on the cell’s repertoire of heat shock factor proteins. Moreover, many desirable cell behaviors (e.g. migration, proliferation, survival/death) cannot be easily controlled at the transcriptional level.

The identification of distinct temperature-responsive proteins, including with functions beyond dimerization, is critical for broad development and application of thermogenetic approaches. Here we introduce a unique thermoresponsive protein module called Melt (<u>Me</u>mbrane localization using temperature), which we derived from the naturally light- and temperature-sensitive BcLOV4 protein ^22^. Melt is a single protein that clusters and binds the plasma membrane at low temperatures but dissociates and declusters upon heating. Using live-cell imaging coupled with custom devices for precise temperature control in 96-well plates^23^, we found that Melt could be toggled between these two states rapidly and reversibly, with observable membrane dissociation and recovery within 10s of minutes. The Melt approach was highly modular, allowing thermal control of diverse processes including EGFR and Ras signaling, TEVp proteolysis, subcellular localization, cytoskeletal rearrangements, and cell death, all through simple end-to-end fusion of the appropriate effectors. We then tuned Melt to increase its switch point temperature above the native 30°C. Such tuning resulted in Melt variants that operated with switch point temperatures between 30-40°C, including ones that bound the membrane at 37°C and fully dissociated at 39°C or 41°C, temperature ranges suitable for downstream application in mammalian tissues. Finally, Melt controlled localized cell death within human cancer xenografts in mice. Thus Melt offers a straightforward, tunable, and broadly applicable platform for endowing thermal control of proteins, cells, and organisms.

## RESULTS

BcLOV4 is a modular optogenetic protein that natively responds to both blue light and temperature ^22,24^ (**Figure 1A**). Light stimulation triggers its clustering and translocation from the cytoplasm to the plasma membrane, where it binds anionic phospholipids ^24,25^. However, its persistence at the membrane requires both continued light and a permissive temperature. At temperatures above 29°C, membrane binding is transient; BcLOV4 binds but then returns to the cytoplasm (**Figure 1A-C**) at a rate that increases with temperature ^22^. Our previous report found that, once dissociated due to elevated temperatures, BcLOV4 remains in the cytoplasm and no longer responds to light stimuli ^22^. However, we found that lowering temperature below the 29°C threshold reversed this inactivation and restored light-dependent membrane localization (**Figure 1C**). Thus, temperature alone could be used to toggle the localization of BcLOV4 given the continued presence of blue light.

**Fig. 1:**
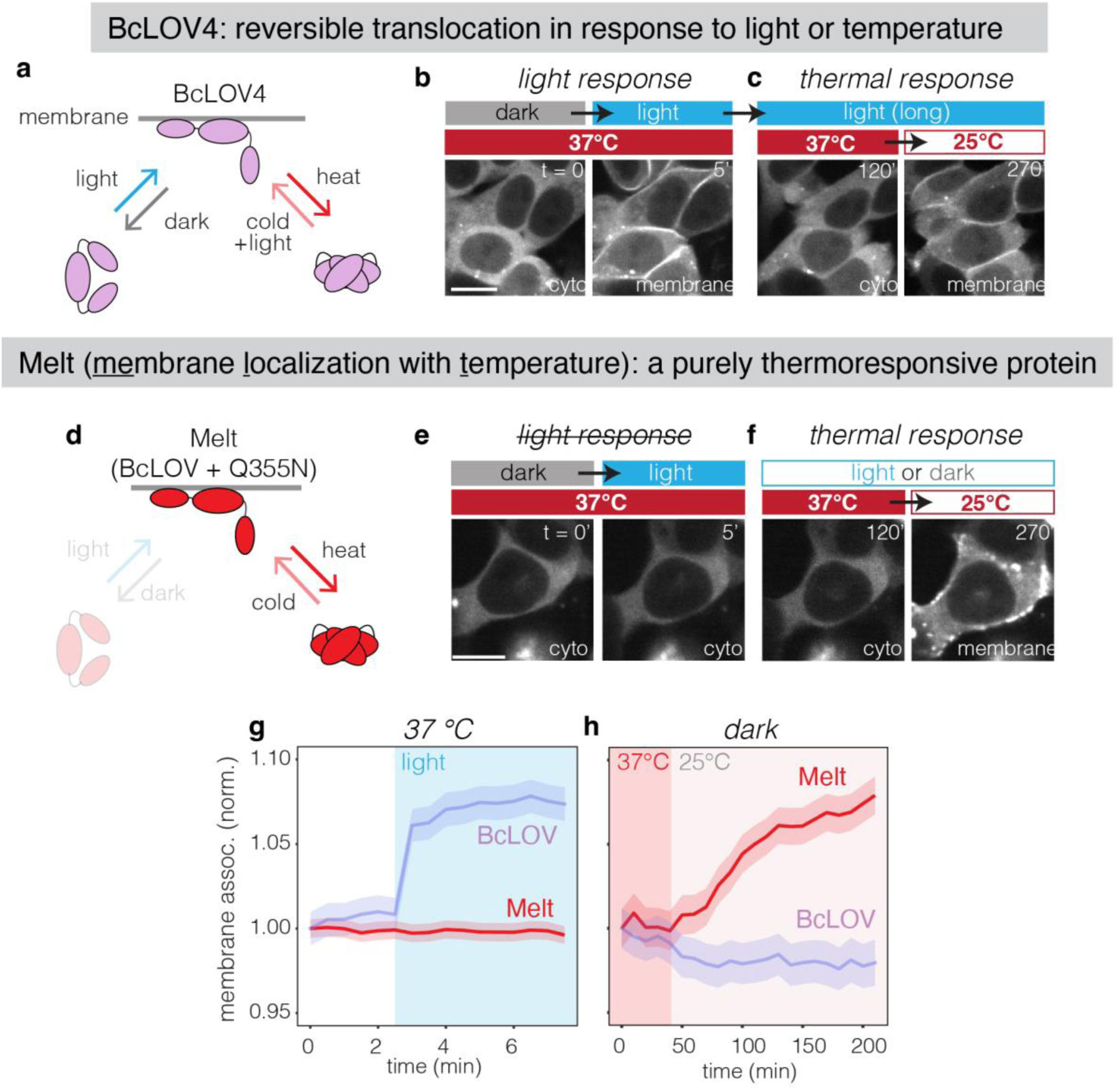
Harnessing BcLOV4 thermosensitivity to generate a purely temperature-inducible protein. A) Schematic of BcLOV4, a naturally light- and temperature-responsive protein. BcLOV4 translocates to the membrane under blue light and reverts to the cytoplasm in the dark. From the membrane-bound (lit) state, elevated temperatures induce dissociation from the membrane, and lower temperatures induce reassociation. B) Representative images showing translocation to the membrane when exposed to blue light in HEK 293T cells. Scale bar = 15 µm. C) Extended illumination at elevated temperatures (2 hr at 37°C, left) causes subsequent disassociation from the membrane, but reversion to lower temperatures (right) allows reassociation with the membrane. D) Schematic of Melt (BcLOV4(Q355N)), which mimics the lit state of BcLOV4. E) Representative images showing that Melt is cytoplasmic at 37°C and does not respond to light (E). However, Melt retains temperature sensitivity and translocates to the membrane upon lowering temperature (F). Scale bar = 15 µm. Comparison of optical (G) and thermal (H) responses of wt BcLOV and Melt. See **Figure S4** for details on quantification. Data represent mean +/- 1 SEM of ∼100 cells. Each construct was normalized to its first timepoint.

We sought to harness this thermal responsiveness to generate a protein actuator that responded only to temperature. We reasoned that a BcLOV4 variant with a point mutation that mimicked the “lit” state would localize to the membrane independent of light status but should retain thermal sensitivity (**Figure 1D**). We thus introduced a Q355N mutation that disrupts the dark-state interaction between the Jα helix and the core of the LOV domain^24,26^, generating a variant that was insensitive to light stimulation (**Figure 1D-G, S1**). In HEK 293T cells at 37°C, BcLOV(Q355N)-mCh was expressed in the cytoplasm. Strikingly, shifting the temperature from 37°C to 25°C triggered an accumulation of the protein at the plasma membrane, where increasing accumulation was observed within minutes and continued over the next three hours (**Figure 1D-H**). In contrast to BcLOV(Q355N), the wt photosensitive BcLOV4 did not accumulate at the membrane in response to temperature in the absence of light (**Figure 1G,H**).

Temperature dependent lipid binding of BcLOV(Q355N)-mCh was also observed when the protein was purified and incubated in a water-in-oil emulsion (**Figure S2**). Thus, BcLOV4(Q355N)—henceforth referred to as Melt (<u>Me</u>mbrane <u>L</u>ocalization using <u>T</u>emperature)—represents a protein whose subcellular localization can be regulated solely by temperature.

Membrane localization of Melt was often accompanied by visible clustering at the membrane, consistent with our prior findings that clustering and membrane-binding are interlinked properties of BcLOV4 ^25,27^ (**Fig 1B,C,F).** Co-expression of Melt-GFP with a CluMPS reporter^28^ and co-immunoprecipitation confirmed that Melt transitioned between a cytoplasmic monomer and a membrane associated oligomer in response to temperature changes (**Figure S3**).

We characterized the thermal response properties of Melt, including how the amplitude and kinetics of membrane dissociation/reassociation varied with time and temperature. To systematically explore this large parameter space, we used the thermoPlate, a device for rapid, programmable heating of 96-well plates^23^. Importantly, the thermoPlate can maintain distinct temperatures in multiple wells simultaneously while also permitting live-cell imaging of the sample using an inverted microscope (**Figure 2A,B**).

**Fig. 2:**
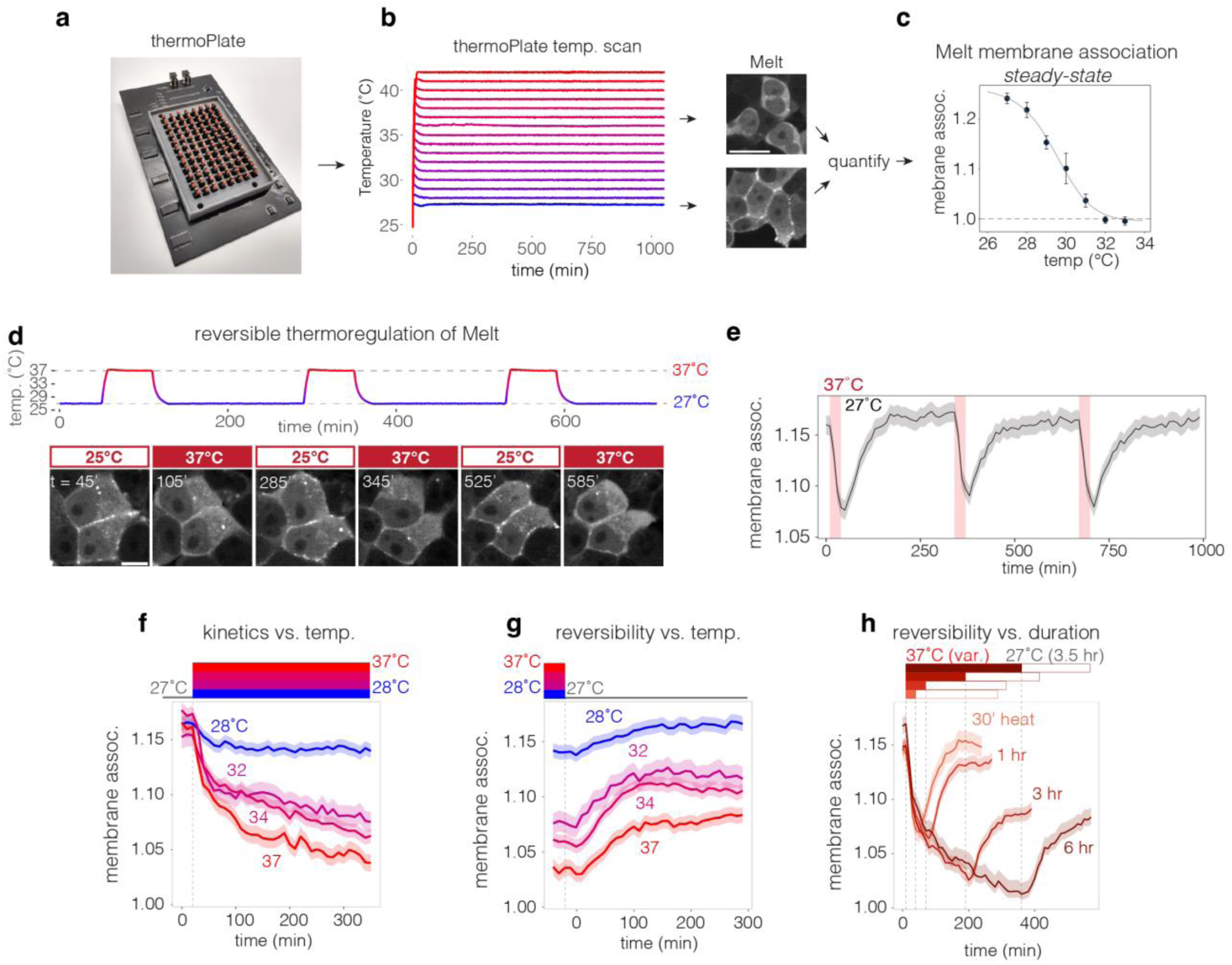
Characterization of Melt membrane association. A) The thermoPlate is a device for thermal control of individual wells in 96-well plate format. B) Heating of 16 individual wells in a 96-well plate with <1°C resolution over 16 hours. Each trace represents the temperature in a single well as recorded by the thermoPlate. Representative images show HEK 293T stably expressing Melt maintained at either 37°C or 28°C via the thermoPlate for 14hr. Scale = 20 µm. C) thermoPlate heating of HEK 293T cells stably expressing Melt allowed measurement of steady-state membrane association (14 hr of heating). Data points represent mean +/- 1 SD of ∼200 cells from each of 3 wells. Dashed line represents membrane association levels of a soluble mCherry. D) Representative images of live-cell images showing Melt membrane binding over multiple cycles of 1 hr at 37°C followed by 3 hr at 27°C. Scale bar = 10 µm. E) Plot of membrane bound Melt while undergoing cycles of 30 min at 37°C followed by 5 hr at 27°C. Traces represent mean +/- 1 SEM of ∼100 cells. F) Kinetics of Melt membrane dissociation when exposed to various temperatures after 24 hr of culture at 27°C. G) Kinetics of Melt membrane reassociation at 27°C after prior exposure to 6 hrs of the indicated temperatures. H) Kinetics of Melt membrane reassociation at 27°C after prior exposure to 37°C for the indicated durations. Each trace in (F-H) represents the mean +/- 1 SEM of ∼1000 cells. Data were collected from HEK 293T cells that stably expressed Melt-mCh.

We first measured steady-state membrane association over a range of temperatures after 14 hrs of heating (**Figure 2C**). Membrane association was maximal at 27°C and minimal at 32°C, and reached 50% of this range at ∼30°C, which we assign as its switch temperature. At temperatures above 32°C, Melt membrane association was undetectable and indistinguishable from that of a soluble mCherry (**Figure S4**). Next, we tested the capacity for dynamic control of Melt. Pulsatile control of temperature between 27°C and 37°C during live cell imaging showed reversible membrane binding and dissociation over multiple cycles (**Figure 2D,E, Supplementary Movie 1**). For full details on membrane binding quantification, see **Figure S4 and Methods.**

We next examined the kinetics of Melt translocation to and from the membrane. Dissociation kinetics increased with higher temperatures (**Figure 2F**). Notably, although steady-state membrane association was unchanged above 32°C (**Figure 2C**), the rate with which Melt reached this steady state level continued to increase with temperature (note the higher decay rate at 34°C and 37°C relative to 32°C, (**Figure 2F**)). Reassociation kinetics depended on the history of thermal stimulation. Samples that were stimulated at higher temperatures showed a lower degree of reversibility (**Figure 2G**). Reversibility was also a function of the duration of prior stimulation. Although dissociation after 30 min of heating at 37°C was fully reversible, longer stimulation led to smaller degrees of reversion (**Figure 2H**). Melt abundance was not affected by high temperature, indicating that incomplete reversion is not due to protein degradation (**Figure S5**). Collectively, these data suggest that Melt is a thermoswitch that operates tunably and reversibly within a 27-32°C range, but whose reversibility is a function of the magnitude of its prior stimulation.

We explored the potential of Melt to control molecular circuits in mammalian cells in response to temperature changes. Recruitment of cargo to/from the membrane is a powerful mode of post-translational control, including for cell signaling ^29^. We first targeted signaling through the Ras-Erk pathway, a central regulator of cell growth and cancer. We generated an end-to-end fusion of Melt to the catalytic domain of the Ras activator SOS2 ^30^, an architecture that previously allowed potent stimulation of Ras signaling using optogenetic BcLOV4 ^22^. We expressed this construct (MeltSOS) in HEK 293T cells and measured Erk activation upon changing temperature from 37°C to 27°C (**Fig 3A**). Active Erk (phospho-Erk, or ppErk) could be observed even within 5 minutes of temperature change to 27°C and continued to rise until its plateau at 30 mins (**Fig 3B,C**). Conversely, shifting temperature from 27°C back to 37°C resulted in measurable signal decrease within 5 min and full decay within 30 mins (**Figure 3B,C**), comparable to the kinetics of thermal inactivation during optogenetic stimulation of BcLOV-SOS ^22^.

**Fig. 3:**
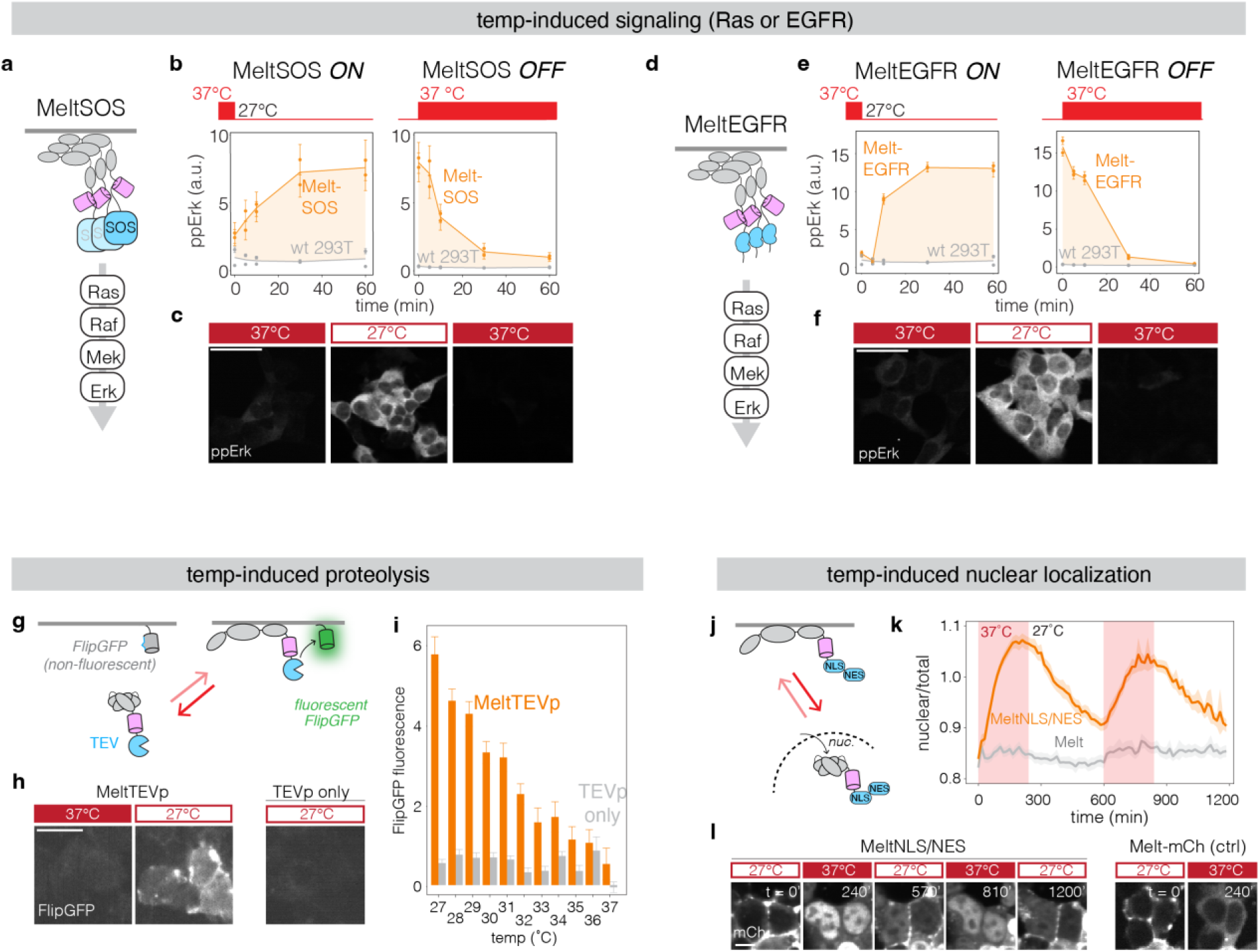
Thermal control over diverse intracellular processes using Melt. A) Schematic of thermal control of Ras-Erk signaling by membrane recruitment of the SOS2 catalytic domain (MeltSOS). B) Thermal activation and inactivation of Ras as assayed by immunofluorescence for activation of the downstream Erk kinase (phospho-Erk, or ppErk). Data points represent the mean +/- 1 SEM of ∼500 cells. C) Representative images of ppErk immunofluorescence from MeltSOS-expressing cells cultured at the indicated temperatures for 24 hours, 1 hour, and 1 hour, respectively. Scale bar = 40 µm. D) Schematic of thermal control of EGFR receptor signaling by membrane recruitment and clustering of the EGFR intracellular domain (MeltEGFR). E) Thermal activation and inactivation of EGFR, assayed through immunofluorescence for ppErk. Each data point represents the mean +/- 1 SEM of ∼500 cells. F) Representative images of ppErk immunofluorescence from MeltEGFR cells cultured at the indicated temperatures for 24 hours, 1 hour, and 1 hour, respectively. Scale bars = 40 µm. G) Schematic of thermal control of proteolysis with MeltTEVp. At low temperatures, MeltTEVp translocates to the membrane where it cleaves a membrane-bound fluorescent reporter of proteolysis (FlipGFP). H) Representative images of FlipGFP fluorescence in cells expressing MeltTEVp or TEVp cultured at 37°C or 27°C for 24 hr. Scale bars = 20 µm. I) Quantification of FlipGFP fluorescence in cells expressing either MeltTEVp or TEVp cultured at the indicated temperature for 24 hours. Each bar represents the mean +/- 1 SEM of ∼1000 cells, normalized between negative and positive controls at each temperature (see **Figure S2** for normalization process). J) Schematic of thermal control of nuclear translocation with MeltNLS/NES. K) Quantification of nuclear localization MeltNLS/NES and Melt-mCh exposed to cycles of 37°C and 27°C. Traces represent the mean +/- 1 SEM of ∼1000 cells. See **Methods** for details on quantification of nuclear localization. L) Representative images of nuclear localization of MeltNLS/NES and Melt-mCh at the temperatures/timepoints found in (K). Scale bar = 10 µm.

Separately, we tested whether we could leverage the clustering of Melt for control of signaling from the receptor level. We generated a fusion of Melt to the intracellular domain of the epidermal growth factor receptor (EGFR) (**Figure 3D**). EGFR is a receptor tyrosine kinase with important roles in development and tumorigenesis and stimulates intracellular signaling through multiple pathways, including Ras-Erk ^31^. Importantly, both membrane recruitment and clustering of the EGFR intracellular domain are required for its activation ^25,32^. In cells expressing MeltEGFR, lowering the temperature from 37°C to 27°C activated strong Erk signaling within 10 minutes, and reversion to 37°C caused signal decay within 5 minutes, with full decay within 30-60 mins (**Figure 3E,F**). Thus, the inducible membrane recruitment and clustering of Melt can be used for rapid, potent, and reversible thermal control of signaling in a modular fashion.

When Melt activates proteins at the membrane, it operates as a heat-OFF system. We next examined whether Melt could also implement a heat-ON system by coupling membrane translocation to negative regulation. Proteases can negatively regulate their targets through protein cleavage in both natural and synthetic systems ^33–35^. We thus tested whether Melt could regulate proteolysis at the membrane. We fused Melt to the viral TEV protease (MeltTEVp) and we measured whether its membrane recruitment could trigger a membrane-associated reporter of TEVp activity, FlipGFP ^36^ (FlipGFP-CAAX). FlipGFP is non-fluorescent until proteolytic cleavage allows proper folding and maturation of the chromophore (**Figure 3G**). Cells that expressed MeltTEVp and FlipGFP-CAAX showed minimal levels of fluorescence when cultured at 37°C, similar to cells that expressed FlipGFP-CAAX and cytoplasmic TEVp or FlipGFP-CAAX alone (**Figure S6)**. However, culturing MeltTEVp cells at lower temperatures for 24 hours increased FlipGFP fluorescence, with fluorescence increasing monotonically with decreasing temperature, whereas cells expressing cytoplasmic TEVp remained at baseline fluorescence (**Figure 3H,I, Figure S6**). Thus, Melt can implement thermal control of proteolysis.

A second way to convert Melt to heat-ON is to regulate its subcellular compartmentalization. Here, the plasma membrane would sequester Melt, and heat would release sequestration and allow translocation to a separate compartment where it could perform a desired function. As a proof of concept, we engineered Melt to regulate nuclear localization by fusing it to sequences that facilitate nuclear import and export (**Figure 3J**). We tested several combinations of nuclear localization sequences (NLS) and nuclear export sequences (NES) to optimize the relative strengths of import and export (**Figure S7**). Melt fused to the SV40 NLS ^37^ and the Stradα NES ^38^ showed strong membrane binding and nuclear exclusion at 27°C and nuclear enrichment when heated to 37°C (**Figure 3K,L, Supplementary Movie 2**). This construct could be dynamically shuttled to and from the nucleus through repeated rounds of heating and cooling. By contrast, Melt without NLS/NES showed no nuclear accumulation upon heating (**Figure 3K,L**). Collectively, our results show that Melt can be applied to control a variety of molecular events, in either heat-ON or heat-OFF configuration, in a straightforward and modular manner.

The utility of Melt in mammals will depend on its ability to induce a strong change in localization in response to temperature, as well as on its ability to switch near mammalian body temperature (∼37°C). We thus sought to tune these properties. To increase the magnitude of membrane translocation, we tested whether short polybasic (PB) peptides could strengthen the electrostatic molecular interactions that mediate BcLOV4 membrane binding (**Figure 4A,B)** ^24,39^. We chose two well-characterized PB domains from the STIM1 and Rit proteins, which can enhance membrane-binding of unrelated proteins ^40^. End-to-end fusions of Melt to the STIM, tandem STIM (STIM2X), or Rit domains all increased the magnitude of membrane binding at 27°C, in increasing order of strength (**Figure 4C,D**). Kinetic analysis showed that PB domains did not change the rate of Melt dissociation, although some changes in reassociation kinetics were observed (**Figure S8**).

**Fig. 4:**
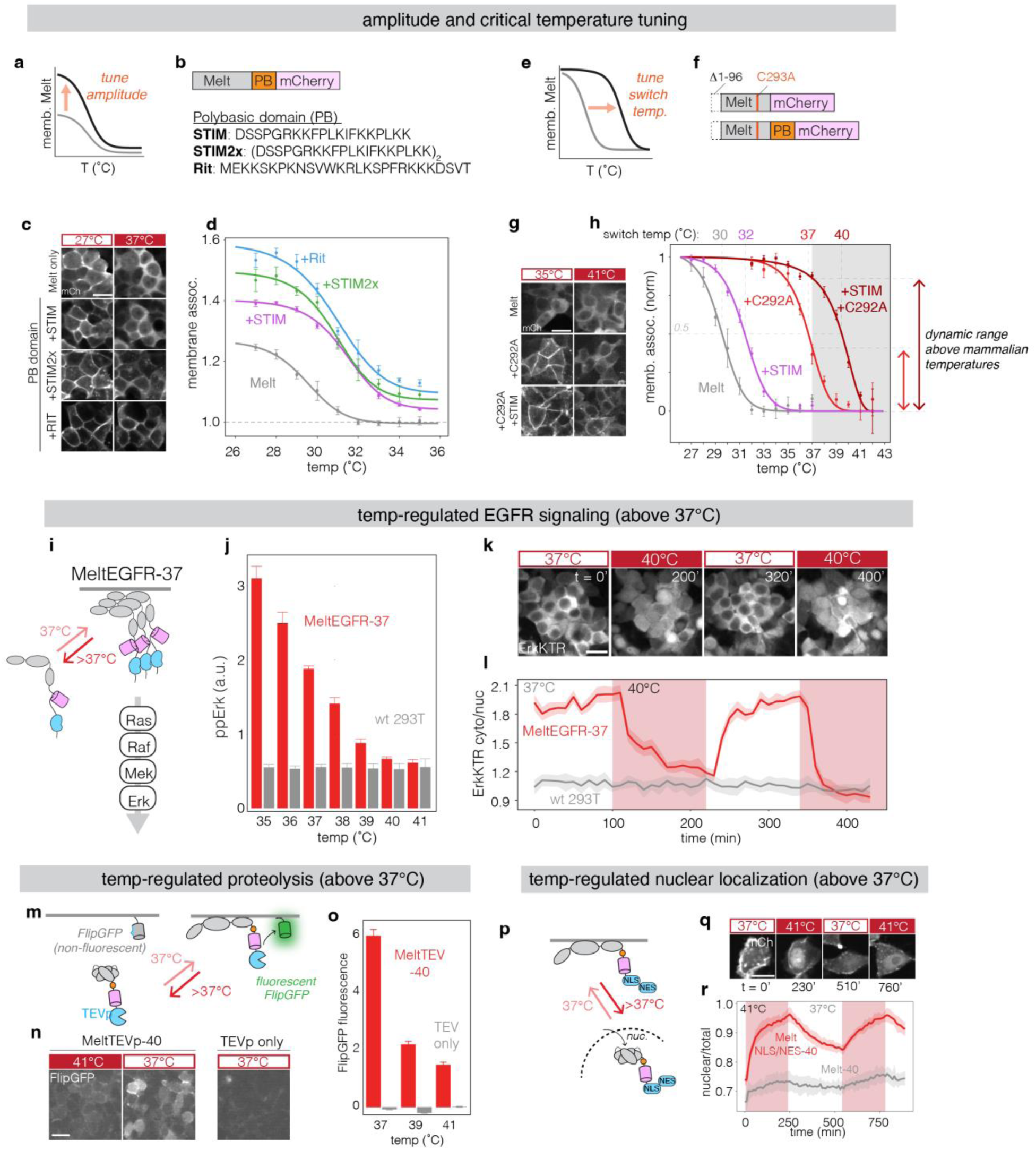
Tuning of Melt membrane binding and thermal switch point allows application of Melt-based tools in mammalian temperature ranges. A) Tuning the amplitude of Melt membrane association. B) Polybasic (PB) domains from the STIM or Rit proteins were fused to Melt to test their ability to increase Melt membrane binding strength. C) Representative images showing stronger membrane binding (higher membrane/cyto ratio) of Melt fused to PBs compared to Melt alone. Melt constructs were stably expressed in HEK 293T cells and are shown after 24 hrs of culture at 27°C and after subsequent heating to 37°C for 6 hrs. Scale bar = 20 µm. D) Quantification of steady-state membrane association of Melt-PB fusions after culture at indicated temperatures for 12 hours. Data represent mean +/- 1 SD of three wells with ∼200 cells quantified per well. Dashed line represents membrane association levels of soluble mCherry. E) Tuning Melt switch-point temperature for use within temperature ranges relevant for mammals, between 37°C and 42°C. F) Schematic of Melt with a C292A mutation with and without STIM PB domain. G) Representative images of membrane localization of Melt, Melt(C292A), or Melt(C292A)+STIM fusion at 35°C for 24 hours and subsequent culture at 41°C for 6 hours. Scale bar = 20 µm. H) Quantification of steady-state membrane binding (14 hrs) of Melt variants between 27 and 42°C. Data represent mean +/- 1 SD of three wells with ∼500 cells quantified per well. Data are normalized between min and max values for each construct. Unnormalized traces can be found in Figure 4D **and Figure S5**. I) Thermal control of EGFR at and above 37°C using Melt-37. J) Immunofluorescence quantification of pathway activation in HEK 293T cells stably expressing MeltEGFR-37. Cells were incubated at indicated temperatures for 75 min before fixation. Bars represent mean +/- 1 SD of three wells with ∼1000 cells quantified per well. K) MeltEGFR-37 activation visualized through the live-cell ErkKTR reporter. Nuclear depletion of ErkKTR indicates Erk activation while nuclear enrichment indicates Erk inactivation. Scale bar = 10 µm. L) Quantification of ErkKTR activity (cyto/nuclear ratio) in HEK 293T cells expressing MeltEGFR-37 or wt cells. Traces represent mean +/- 1 SD of ∼15 cells per condition. M) Control of proteolysis at mammalian temperatures with MeltTEVp-40. N) Representative images of FlipGFP signal in cells expressing MeltTEVp-40 or TEVp after incubation at the indicated temperatures for 24 hours. Scale bar represents 10 µm. O) Quantification of FlipGFP signal in fixed cells expressing MeltTEVp-40 or TEVp cultured at the indicated temperatures for 24 hours. Each bar represents the mean +/- 1 SEM of ∼1000 cells. Y-axis represents mean fluorescence subtracted by the signal of TEVp-negative cells. P) Control of nuclear translocation at mammalian temperatures with MeltNLS/NES-40. Q) Representative images of nuclear translocation. Scale bar = 20 µm. R) Quantification of nuclear localization of MeltNLS/NES-40 or Melt-40-mCh after exposure to cycles of 37°C and 41°C (red) in HEK 293T cells. Traces represent the mean +/- 1 SEM of ∼1000 cells.

Although PB domains provided a large increase in steady-state membrane binding at 27°C, they provided only a mild increase in thermal switch point to ∼32°C, only 1-2 degrees higher than the original Melt (**Figure 4D).** We achieved a more substantial increase through the fortuitous discovery that the C292 residue plays an important role in defining the Melt thermal response. In wt BcLOV4, C292 is thought to form a light-dependent bond with a flavin mononucleotide cofactor that underlies the BcLOV4 photoresponse ^24^. Although Melt translocation does not respond to light (**Figure 1G, S1)**, introduction of a C292A mutation dramatically increased its membrane association not only at 27°C, but also at 37°C where the original Melt was fully dissociated **(Figure 4E-H, Figure S9**). As before, addition of the STIM PB domain further increased membrane association strength at these higher temperatures.

Importantly, both C292A variants retained temperature sensitivity and dissociated from the membrane at 41-42°C, with a thermal switch point of 36.5 and 39.5°C for the C292A and C292A/STIM variants, respectively (**Figure 4H, Figure S9,10**). Because these Melt variants can exist in one state at 37°C and another at 41/42°C, they are thus both potentially suitable for heat activation within mammalian tissues, with distinct levels of membrane binding and dynamic range that could each be optimal for certain applications. These variants also included a truncation of 96 amino acids from the N-terminal of BcLOV4, which we found expendable, consistent with previous results ^24^. Collectively, our work presents four Melt variants with a range of thermal switch points between 30°C and 40°C, covering temperatures suitable for actuation in cells from a broad range of species. We adopted a nomenclature for these variants that reflects these switch points: Melt-30, Melt-32, Melt-37, and Melt-40.

We tested the ability of the higher switch-point Melt variants to actuate post-translational events between 37 and 42°C. MeltEGFR driven by Melt-37 showed strong Erk activation at 37°C and only baseline levels at 40-41°C (**Figure 4I,J**). Erk activity could be stimulated repeatedly over multiple heating/cooling cycles as indicated by the ErkKTR biosensor, which translocates from the nucleus to the cytoplasm upon Erk activation (**Figure 4K,L, Supplementary Movie 3**) ^41^. MeltSOS-37 could also stimulate Erk activity but only at <∼37°C, potentially reflecting a requirement for higher levels of membrane translocation relative to MeltEGFR ^25^ (**Figure S11)**.

Melt-37/40 could also regulate proteolysis and protein translocation. Melt-40 fused to TEVp showed strong proteolysis and FlipGFP activation at 37°C, with markedly reduced activity at 41°C (**Figure 4M-O**). Melt-37 also regulated proteolysis but only induced fluorescence at or below 35°C, and fluorescence fell to near baseline at 37°C (**Figure S12**). These results further highlight that although the general thermal response properties are dictated by the specific Melt variant, the precise thermal switch point of the downstream process can be influenced by the specific fusion partner or the downstream process itself. Melt-40 also regulated membrane-to-nuclear translocation within the well-tolerated 37-41°C temperature range (**Figure 4P**). Fusion to a C-terminal SV40 NLS and Stradα NES allowed strong membrane sequestration at 37°C, and fluorescence became enriched in the nucleus upon heating to 41°C (**Figure 4Q,R**). As before, translocation was partially reversible on the timescales tested and could be cycled through repeated rounds of heating and cooling (**Figure 4Q,R, Supplementary Movie 4**).

We then asked whether Melt variants could be used to regulate cellular-level behaviors at and above 37°C. We first sought to control cell shape changes through the control of actin polymerization. We fused Melt-37 to the DH-PH domain of Intersectin1 (MeltITSN1-37), an activator of the Rho GTPase Cdc42 that has previously been actuated through optogenetic recruitment ^42^, including with BcLOV4 ^43,44^ (**Figure 5A**). When cooled from 41°C to 37°C, HEK 293T cells expressing MeltITSN1 showed rapid and dramatic expansion of lamellipodia and cell size, consistent with Cdc42 activation ^45^ (**Figure 5B**). Changes in cell shape could be reversed and re-stimulated over multiple cycles of cooling and heating (**Figure 5C**), showing similar magnitude of shape change in each round (**Figure 5D, S13, Supplementary Movie 5**). By comparison, temperature changes had no effect on cell shape in cells that expressed Melt-37 without the ITSN1 DH-PH domain.

**Fig. 5:**
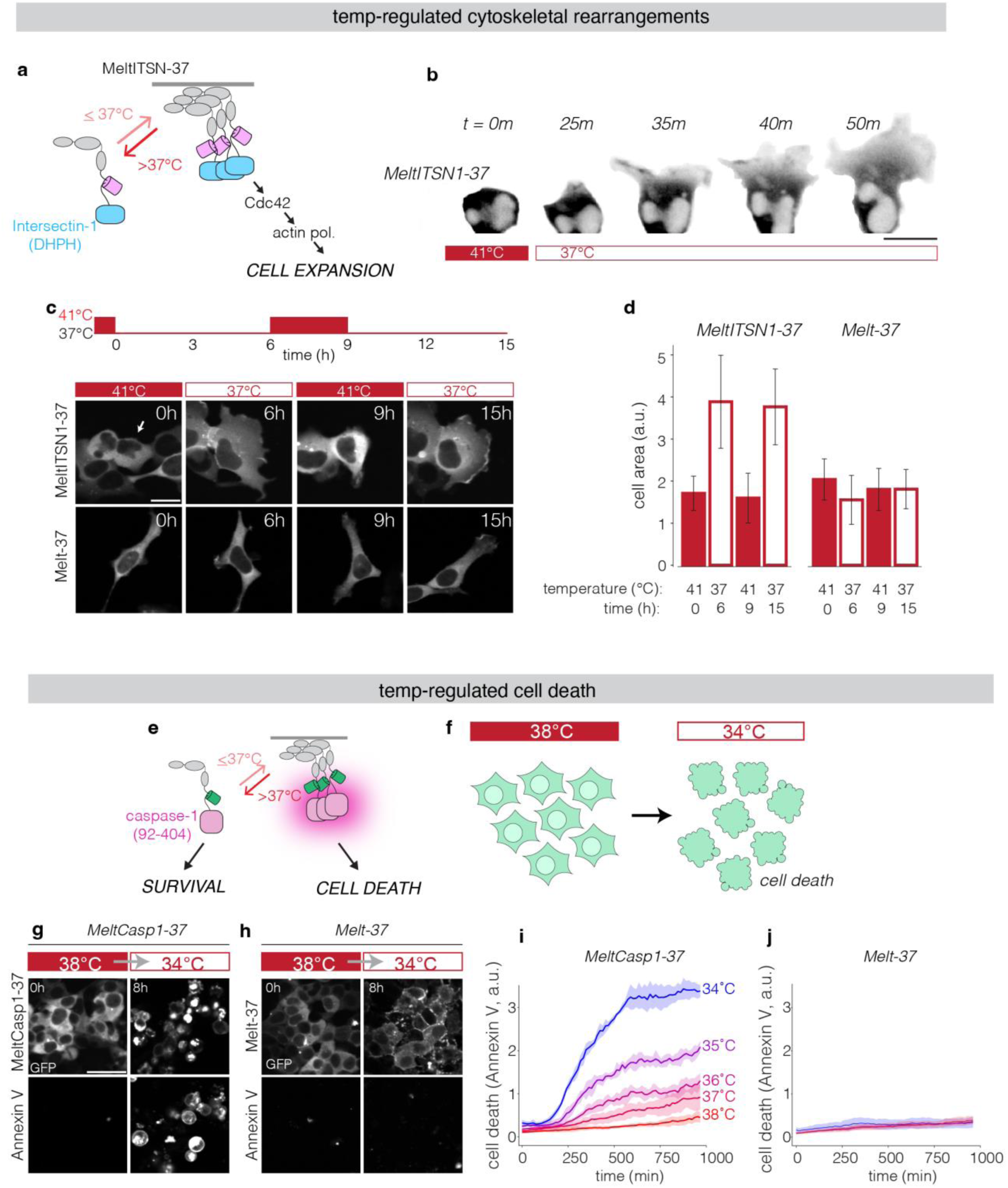
Thermal regulation of cell fate using Melt. A) Control of Cdc42 activity and cell shape through recruitment of the DHPH domain of ITSN1 to the membrane. B) Representative images of cell shape changes in response to temperature control in a HEK 293T cell transiently expressing MeltITSN1-37. Upon reduction of temperature from 41°C to 37°C, cells show rapid formation of membrane extensions and dramatic increase in size. Scale bars = 20 µm. C) Cell shape changes are reversible and repeatable over several hours of stimulation. Representative images of HEK 293T cells transiently transfected with MeltITSN1-37, cultured at 41°C and exposed to multiple rounds of heating and cooling at the times and temperatures indicated. Scale bars = 20 µm. D) Quantification of cell area of cells expressing either MeltITSN1-37 or Melt-37 after repeated cooling and heating. Bars represent the average cell size of 15 cells +/- 1 SD. E) Thermal control of cell death through regulation of caspase-1 clustering (MeltCasp1-37).F) MeltCasp1-37 induces cell death upon lowering temperature below 37°C. G) Representative images of cells expressing MeltCasp1-37 (G) or Melt-37 (H) before and after exposure to 34°C for 8 hours after culture at 38°C for 24 hours. Bottom panels of (G,H) show Annexin V-647 staining, which indicates cell death. Scale bars = 40 µm. I) Quantification of Annexin V intensity in MeltCasp1-37 and Melt-37 cells over time at the indicated temperature after prior culture at 38°C for 24 hours. Plots represent the mean +/- SEM of per-image Annexin V fluorescence divided by total GFP fluorescence (to account for cell density) across 4 images. See **Methods** for quantification details. All images/data in this figure were collected using transient expression of Melt constructs in HEK 293T cells.

As a second example, we asked if Melt could be used for thermal control of cell death. Cell death can be achieved by regulated clustering of effector domains of caspase proteins ^46^. We fused Melt-37 to the effector domain of caspase-1 (MeltCasp1-37, **Figure 5E**), and we measured cell death upon changes in temperature (**Figure 5F**). Cells expressing MeltCasp1-37 appeared unperturbed at 38°C, a further indicator that Melt is monomeric at elevated temperatures, as even dimers of the caspase-1 domain cause cell death (**Figure S14**). By contrast, lowering of temperature to 34°C led to morphological changes within minutes, followed within hours by blebbing and cell death, indicated by both morphology and Annexin V staining (**Figure 5G,H, Supplementary Movie 6**). ThermoPlate scanning coupled with live cell imaging of Annexin V revealed cell death induction even when shifting temperature by only 1°C (from 38°C-37°C), and the magnitude of cell death increased with larger temperature shifts (**Figure 5I,J**). No death was measured in cells expressing Melt-37 without the caspase effector.

A potential concern for using heat as a cellular stimulus is that heat is a known stressor and could adversely affect cell functions. However, we observed no molecular or functional effects of either the short- or long-term heat profiles used throughout our studies in mammalian cells. Stress granules (SGs), a known consequence of heat-stress^47,48^, were not observed at or below 41°C in HEK 293T cells, the operating temperatures for the highest switch-point Melt variants (**Figure S15A,B)**. By contrast, SGs could be detected at 42°C in ∼1-5% of cells, and at 43°C all cells showed strong SG formation. Of note, many existing strategies for thermal induction are typically stimulated with 42°C^4,13,17,18^, at the cusp of this non-linear heat-induced SG response (**Figure S15B**). We also measured cell proliferation to investigate potential integration of low-level heat stress during multi-hour heating (**Figure S15C,D)**. No differences in proliferation were observed when cells were cultured for 24 hrs at temperatures up to 41°C, the highest temperature required to stimulate our Melt variants. Growth defects appeared only at 42°C and above.

Thermogenetics offers the exciting potential for remote, dynamic, and spatially-resolved control of cells within opaque tissues that are inaccessible to alternative dynamic stimuli like light. To test this premise, we developed a tissue-mimicking phantom to model various tissue depths^49^ (**Figure S16A)**, and we tested the ability for light or temperature to stimulate clustering of the caspase1 fragment. While direct illumination of a light-sensitive caspase-1^46^ resulted in strong killing, illumination through 2mm of the phantom reduced killing by ∼75%, and killing was undetectable at increased thicknesses (**Fig S16B**). By contrast, MeltCasp1-37 induced cell death independent of phantom thickness at 34°C.

Finally, we asked whether Melt could control cell behavior in animals in a spatiotemporally defined manner by testing its ability to induce cell death in mouse xenografts of human cancer cells. H3122 lung cancer cells expressing MeltCasp1-37 and firefly luciferase rapidly underwent cell death in < 3hrs after cooling from 37-25°C in culture (**Figure 6A**). We then injected these cells into both flanks of immunodeficient NSG mice and, 48 hr after injection, we cooled the tumor on one flank while leaving the contralateral tumor untreated (**Figure 6B**).

**Fig 6.**
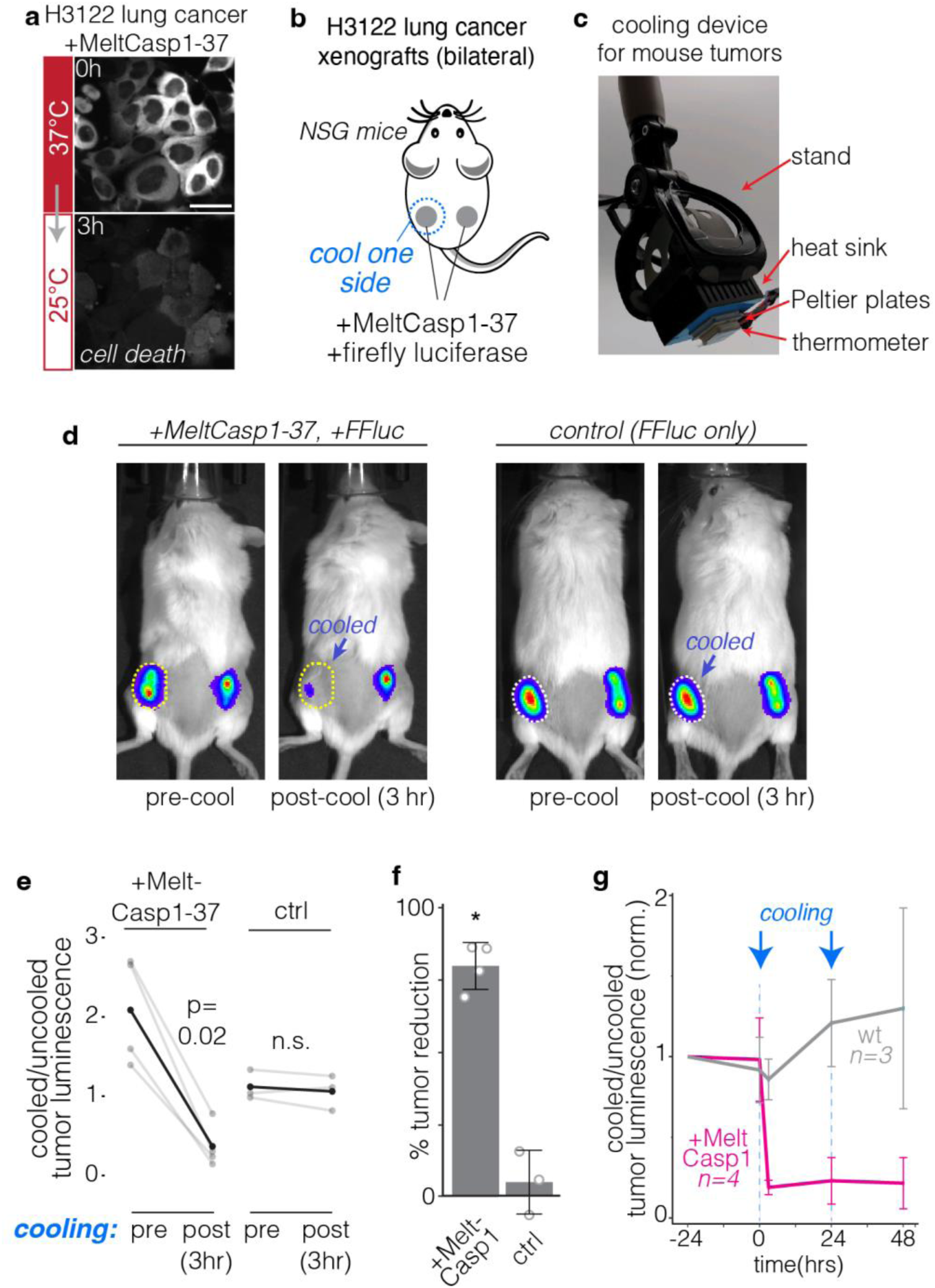
Thermal control of Melt and cell fate in animal models. A) H3122 cancer cells expressing MeltCasp1-37 show rapid cell death within 3 hrs after cooling. Scale bar represents 40µm. B) Bilateral tumor model to test spatiotemporal control of MeltCasp1-37 *in vivo*. Mice were injected on both flanks with H3122 cells expressing MeltCasp1-37 and firefly luciferase or luciferase only (control). 48 hrs post injection, cooling was applied locally to one flank. C) Device for programmable cooling of xenografts. A Peltier element cools the outward-facing surface, which provides localized topical cooling when applied to the mouse. A thermistor allows real-time monitoring and feedback control of temperature. D) Representative images of mice before and after cooling. Cold treated tumors showed dramatic reduction in luciferase signals relative to uncooled tumors, but only for tumors expressing MeltCasp1-37. Cooling protocol: 45 minutes of 5°C followed by 45 minutes of 15°C. E) Quantification of (D). Light grey: individual mice. Black: mean. Significance determined by a one-sided Wilcoxon signed-rank test. F) Tumor reduction was obtained by calculating the change in luminescence ratio (cooled/uncooled) as a result of cooling. N = 3-4 mice, bars = mean +/- SD. p = 0.028 by one-sided Mann-Whitney test. F) Relative luminescence of cooled vs uncooled tumors over multiple days, with treatment repeated at 0 and 24 hrs. Traces represent the mean +/- 1 SD of 3-4. Values normalized to the first day of imaging.

Cooling (45 min at 5°C followed by 45 min at 15°C) was performed by topical application of a custom thermoelectric cooling device that maintained programmable feedback-controlled temperature (**Figure 6C, Figure S17**). Luciferase imaging revealed ∼80% reduction of tumor cells in the cooled flank relative to the uncooled flank only 3 hr after cooling. No reduction of tumor cells was observed in xenografts lacking MeltCasp1-37 (**Figure 6D,E,F**). Cooling over subsequent days gave no further reduction in luciferase signal, suggesting that the initial cooling maximally eliminated cells (**Figure 6G**). Thus, Melt can control cell behavior in mammals in a spatiotemporally defined manner using a non-invasive temperature stimulus.

## DISCUSSION

Here we have described a modular and tunable protein that permits thermal control over a range of molecular and cell-level behaviors. By locking the naturally light- and temperature-sensitive BcLOV4 into its “lit” state, we generated the purely thermoresponsive Melt whose membrane association and clustering can be regulated with a small temperature change (<4°C). Tuning this thermal response further allowed us to generate multiple variants (Melt-30/32/37/40) whose activation switch points could be shifted within the 30-40°C range. These variants allowed temperature-inducible control of signaling, proteolysis, and subcellular localization, including between 37°C-42°C, a critical range for thermal control within mammals. Finally, we showed that Melt can provide thermal control over cell and tissue-level behaviors by changing cell size/shape and cell death, both *in vitro* and *in vivo*.

Our engineering efforts provide insight into how the wt BcLOV4 protein senses both light and temperature. Successful isolation of the BcLOV4 thermal response from its light response confirms the distinct molecular nature of these two behaviors, as previously speculated ^22^. At the same time, the light and temperature responses are intertwined, since mutation of the C292 residue in the LOV domain, which mediates photo-responsiveness, dramatically shifted the thermal switch point of Melt (**Figure 4E**). Nevertheless, the molecular mechanism of thermosensing remains unclear. One possibility is that higher temperatures generate a new intramolecular interaction that occludes the membrane-binding interface of Melt/BcLOV4. This could be achieved either directly through an interaction interface that strengthens at higher temperature, or via partial unfolding of a domain that reveals a new binding interface. Future mechanistic studies will provide clarity here and will allow optimization of Melt properties including speed of response and degree of reversibility, and will shed light on how the photosensing and thermosensing elements of BcLOV4 interact. These latter studies will additionally provide insight for how to engineer novel multi-input proteins that can perform complex logic in response to user-defined stimuli.

Multiplexed control of sample temperature allowed us to systematically characterize new Melt variants, ultimately resulting in variants with switch-points ranging from 30-40°C. Because optogenetic BcLOV4 works in mammalian cells but also in systems that are cultured at lower temperatures like yeast, flies, zebrafish, and ciona^22,24,43,50–52^, we anticipate that all Melt variants will find use across these and similar settings. Our work also highlights the utility of having multiple variants in hand to optimize specific downstream applications. We found on multiple occasions that the precise thermal response profiles depended not only on the specific Melt variant but also on both the effector and downstream process under control, thus requiring empirical validation for each use case and biological context. Optimization can be performed by testing other Melt variants, or by generating new ones through additional mutations or modifications (e.g. polybasic domains) similar to the ones we describe.

While the benefits of penetrant, spatiotemporally precise control could in principle be achieved using other stimuli like magnetic fields or sound waves, these approaches are limited by the lack of biomolecules that respond to these inputs. In this respect, thermal control is a more practical and tractable approach. Still, there remain surprisingly few strategies for engineering thermally-controllable protein systems.

Melt dramatically expands the range of molecular and cellular events that can be controlled by temperature and, in mammalian cells, allows thermal control with lower potential for heat stress relative to the few existing approaches. Melt provides an orthogonal input control that can be used in conjunction with—or instead of—existing technologies based on light or chemicals, and it affords unique potential for actuation of proteins and cells in animals, opening exciting avenues across biotechnology and biomedicine.

## Supporting information

Move S1

## Acknowledgements

We thank Erin Berlew and Brian Chow for helpful discussions on BcLOV4 activity and for plasmids encoding BcLOV(Q355N) and BcLOV-ITSN1, and Alex Hughes and Matthew Good for helpful comments on the manuscript. We also thank the Penn Cytomics and Cell Sorting Shared Resource Laboratory for assistance with cell sorting. This work was supported by funding from the National Institutes of Health (R35GM138211 for L.J.B), the National Science Foundation (Graduate Research Fellowship Program to W.B., CAREER 2145699 to L.J.B.), and the Penn Center for Precision Engineering for Health. Cell sorting was performed on a BD FACSAria Fusion that was obtained through NIH S10 1S10OD026986.

## Code and Data Availability

All data and code found in this manuscript can be accessed at https://rb.gy/1k7tc. All raw images are available on request. All unique biological materials are available upon request.

## Author Contributions

W.B. and L.J.B. conceived the study to generate Melt and downstream applications. W.B. generated Melt and its integration into molecular circuits. Z.H. discovered and characterized thermostable Melt variants, which were then integrated into circuits by Z.H. and W.B. W.B. and P.I. developed and validated the thermoPlate. D.W. and T.R.M. validated cluster-induced cell killing. W.B., Z.H., and P.I. performed and analyzed all experiments. L.J.B. supervised the work. W.B., Z.H., and L.J.B. wrote the manuscript and made figures, with editing from all authors.

## List of Supplementary Materials

Materials and Methods. Supplementary Figures 1-17. Supplementary Movie Captions 1-6.

## METHODS

### Cell Culture

Lenti-X HEK 293T cells were maintained in 10% fetal bovine serum (FBS) and 1% penicillin/streptomycin (P/S) in DMEM. (Lenti-X HEK 293T: Takarabio 632180). Cell lines were not verified after purchase. Cells were not cultured in proximity to commonly misidentified cell lines.

### Plasmid design and assembly

Constructs for stable transduction and transient transfection were cloned into the pHR lentiviral backbone with a CMV promoter driving the gene of interest. Melt mutations were introduced to WT BcLOV4 (Provided by Brian Chow) (Addgene Plasmid #114595) via whole backbone PCR using primers containing the target mutation. Mutations were introduced using the same primers on BcLOV4-ITSN1 (Provided by Brian Chow) (Addgene #174509) to generate MeltITSN1-37. Melt-PB fusions were generated via whole backbone PCR using primers containing PB coding sequences (**Figure 2B**). PCR products were circularized via ligation (New England Biolabs). For Melt-effector fusions, the pHR backbone was linearized using MluI and NotI restriction sites. Melt, TEVp (Addgene Plasmid #8827), EGFR (sourced from Opto-hEGFR, which was a kind gift from Dr. Harold Janovjak), SOS ^22^, and Caspase-1 (Provided by Peter Broz) ^46^ were generated via PCR and inserted into the pHR backbone via HiFi cloning mix (New England Biolabs). All Melt37/40-Effector fusions were generated by amplifying Melt37/40 with primers that amplified the region downstream of a.a.96 such that the final Melt variants contained a a.a.1-96 deletion. NLS/NES insertions were generated via backbone PCRs with NLS/NES sequences (**Figure S3**) incorporated into the primers. To construct FlipGFP-BFP-CAAX, the two fragments of FlipGFP B1-9 and B10-E5-B11-TEVcs-K5 were amplified from Addgene Plasmid #124429 via PCR. tagBFP ^22^ was amplified using primers containing a CAAX membrane binding sequence. These fragments were assembled in the linearized PHR backbone via HiFi cloning mix in the order B1-9-P2A-B10-E5-B11-TEVcs-K5-tagBFP-CAAX. In order to reduce affinity of TEVp for the TEV cut site (cs) and lower basal proteolysis, the canonical cut site ENLYFQS was mutated to ENLYFQL ^53^ via whole backbone PCR using primers harboring the mutation. GFP-CAAX was generated via PCR of eGFP using primers containing the CAAX sequence and cloned into the linearized viral backbone using HiFi cloning mix.

### Plasmid transfection

HEK 293T cells were transfected using the calcium phosphate method, as follows: Per 1 mL of media of the cell culture to be transfected, 50 µL of 2x HeBS^28,29^ buffer, 1 µg of each DNA construct, and H_2_O up to 94 µL was mixed. 6 µL of 2.5mM CaCl_2_ was added after mixing of initial components, incubated for 1:45 minutes at room temperature, and added directly to cell culture.

### Lentiviral packaging and cell line generation

Lentivirus was packaged by cotransfecting the pHR transfer vector, pCMV-dR8.91 (Addgene, catalog number 12263), and pMD2.G (Addgene, catalog number 12259) into Lenti-X HEK293T. Briefly, cells were seeded one day prior to transfection at a concentration of 350,000 cells/mL in a 6-well plate. Plasmids were transfected using the calcium phosphate method. Media was removed one day post-transfection and replaced with fresh media. Two days post-transfection, media containing virus was collected and centrifuged at 800 x g for 3 minutes. The supernatant was passed through a 0.45 µm filter. 500 µL of filtered virus solution was added to 700,000 HEK293T cells seeded in a 6-well plate. Cells were expanded over multiple passages, and successfully transduced cells were enriched through fluorescence activated cell sorting (Aria Fusion).

### Preparation of cells for plate-based experiments

All experiments were carried out in Cellvis 96 well plates (#P96-1.5P). Briefly, wells were coated with 50uL of MilliporeSigma™ Chemicon™ Human Plasma Fibronectin Purified Protein fibronectin solution diluted 100x in PBS and were incubated at 37 °C for 30 min. HEK 293T cells were seeded in wells at a density of 35,000 cells/well in 100 µL and were spun down at 20 x g for 1 minute. In experiments requiring starvation (for all experiments involving SOS and EGFR constructs), after 24 hr, cells were starved by performing 7 80% washes with starvation media (DMEM + 1% P/S). Experiments were performed after 3 hr of starvation.

### Fixing and Immunofluorescence staining

Immediately following the completion of a temperature stimulation protocol, 16% paraformaldehyde (PFA) was added to each well to a final concentration of 4%, and cells were incubated in PFA for 10 min. For immunofluorescence staining, cells were then permeabilized with 100 µL phosphate buffered saline (PBS) + 0.1% Triton-X for 10 min. Cells were then further permeabilized with ice cold methanol for 10 min. After permeabilization, cells were blocked with 1% BSA at room temperature for 30 min. Primary antibody was diluted in PBS + 1% BSA according to the manufacturer’s recommendation for immunofluorescence (phospho-p44/42 MAPK (Erk1/2) (Thr202/Tyr204), Cell Signaling #4370, 1:400 dilution; phospho-Rb (Ser807/811) Cell Signaling #9308, 1:800 dilution; Anti-Human G3BP1, BD Biosciences #611126, 1:500 dilution). Wells were incubated with 50 µL of antibody dilution for 2 hr at room temperature (RT), after which primary antibody was removed and samples underwent five washes in PBS + 0.1% TWEEN-20 (PBS-T). Cells were then incubated with secondary antibody (Jackson Immunoresearch Alexa Fluor® 488 AffiniPure Goat Anti-Rabbit IgG (H+L) or Invitrogen Goat anti-Mouse IgG (H+L) Cross-Adsorbed Secondary Antibody, DyLight™ 650) and DAPI (ThermoFisher, #D1306, 300 nM) in PBS-T + 0.1% BSA for 1 hour at RT. Secondary antibody was removed, samples underwent 5 washes with PBS-T. Samples were imaged in PBS-T.

### Imaging

#### Live-cell imaging

Live-cell imaging was performed using a Nikon Ti2-E microscope equipped with a Yokagawa CSU-W1 spinning disk, 405/488/561/640 nm laser lines, an sCMOS camera (Photometrics), a motorized stage, and an environmental chamber (Okolabs). HEK 293Ts expressing the construct of interest were imaged with a 20X or 40X objective at variable temperatures and 5% CO_2_. Optogenetic BcLOV4 was stimulated using a 488nm laser.

#### High content fixed-cell imaging

Fixed samples were imaged using a Nikon Ti2E epifluorescence microscope equipped with DAPI/FITC/Texas Red/Cy5 filter cubes, a SOLA SEII 365 LED light source, and motorized stage. High content imaging was performed using the Nikon Elements AR software. Image focus was ensured using image-based focusing in the DAPI channel.

### Image processing and analysis

#### Immunofluorescence quantification

Images were processed using Cell Profiler. Cells were segmented using the DAPI channel, and cytoplasm was identified using a 5 pixel ring around the nucleus. Nuclear and cytoplasmic fluorescence values were then exported and analyzed using R (https://cran.r-project.org/) and R-Studio (https://rstudio.com/). Data was processed and visualized using the tidyR ^54^ and ggplot2 ^55^ packages.

#### Membrane recruitment

Membrane localization was quantified using the MorphoLibJ plugin for ImageJ ^56^. Briefly, MorphoLibJ was used to segment single cells based on a constitutively membrane bound GFP-CAAX marker. The resulting segmentation was imported into Cell Profiler and was used to quantify the mean mCherry (fused to the protein of interest) localized to the membrane as well as mean mCh per cell (**Figure S4**). Mean mCh and membrane-localized mCh intensity was recorded and further processed in R. Differences in expression levels were corrected for by dividing the mean membrane intensity of mCh by mean cell mCh. Membrane binding data was then normalized such that minimum membrane binding was represented as 1.0 to match the membrane binding levels of a cytoplasmic mCh, as detailed in **Figure S4**.

#### FlipGFP Quantification

Cells expressing membrane bound FlipGFP-CAAX and the indicated TEVp construct were grown at the indicated temperature and fixed in 4% PFA after 24 hours. FlipGFP was tethered to the membrane via a Blue Fluorescent Protein (TagBFP)-CAAX fusion. BFP-CAAX remained tethered to the membrane before and after proteolysis and thus could be used as a membrane marker. This marker was used to segment single cells using the same workflow used for membrane recruitment quantification. Single cell GFP levels were quantified using Cell Profiler and used as an indicator of relative levels of proteolysis.

#### Nuclear Localization

To quantify nuclear localization of a protein of interest, cells expressing a GFP-CAAX membrane marker (see above) were transfected with an H2B-iRFP nuclear marker. The above workflow was used to segment individual cells based on the membrane marker. This segmentation was imported to CellProfiler, which was also used to segment nuclei based on iRFP imaging. Each nucleus was then assigned to a parent cell. Nuclei were assigned to a cell if >90% of the nucleus object was contained by the cell object. Membrane segmented cells that contained no nuclei objects or nuclei that were not within a parent cell were eliminated from quantification. Finally, nuclear to total cell mCherry (used as a marker fused to the protein of interest) was calculated and recorded for each cell.

#### Annexin Staining and Quantification

Annexin V-647 (Invitrogen A23204) was added to 100 µL of cell culture at a 1:100 final dilution. A final concentration of 1 mM CaCl_2_ was also added to each well to allow Annexin V cell labeling. Cell media was removed and replaced with Annexin V media 30 min prior to imaging. To quantify Annexin V, images of cells expressing MeltCasp1-37 or Melt-37 both with a GFP fusion were used to create GFP masks using CellProfiler’s threshold function. Annexin images were masked for GFP positive pixels. The total masked Annexin image intensity was recorded and normalized by the number of GFP positive pixels (cell area per image) in each image.

#### Cell Area Quantification

Cell area was measured semi-manually. Images of cells expressing MeltITSN1-37 and Melt-37 were imaged and resulting images were thresholded in ImageJ such that cell positive pixels were set to 1 and background pixels were set to 0. Cells were manually chosen for quantification and regions containing the cell of interest were drawn by hand. Measuring integrated pixel intensity of these regions gave rise to the number of cell positive pixels in that region which was used as a metric of total cell area. For further explanation, see **Figure S9**.

### Curve fitting

Data points for Melt variant equilibrium membrane binding at various temperatures were fit to the Hill Equation (Eq.1). MATLAB was used to minimize the error between the sigmoid function and each data point. The characteristic function used for fitting was:

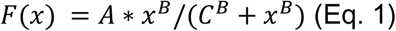

A, B, and C were used as the adjusted parameters. These curves are displayed in **Figure 2E, 4D, and 4H** with datapoints overlaid. The associated code can be found in this manuscript’s code repository (https://rb.gy/1k7tc).

### Protein purification

HisTag-GB1-Melt-mCh-HisTag was transformed in *E. coli* strain BL21 for protein production. Bacteria was inoculated into 5 ml fresh LB media for overnight growth at 37°C. 1:100 dilution was performed to amplify the culture in 500 ml until OD600 reached 0.4-0.8 at 37°C. Then IPTG was added to 0.5 mM for protein production at room temperature (22°C) for 24-36 hours. Bacteria were then pelleted and frozen at −20°C for 20 minutes and then lysed with lysis buffer (50 mM Na_2_HPO_4_, 500 mM NaCl, 0.5% Triton-X-100 and protease inhibitor at pH 6.5) and sonicated. The following steps were performed under 4°C. The sample was then sedimented by centrifugation (15400 x g for 60 min in 15 mL tubes), and the supernatant was loaded on columns containing nickel resins (TaKaRa #635506) and mixed at 4°C for 20 min. The columns were washed with 2 mL of 10 mM imidazole dissolved in wash buffer (50 mM Na2HPO4, 500 mM NaCl, 10% glycerol, and protease inhibitor at pH 6.5), 2 ml PBS, 500 µL100 mM imidazole dissolved in the same wash buffer. Finally, 500 µL elution buffer (50 mM Na2HPO4, 500 mM NaCl, 10% glycerol, 500 mM imidazole, and protease inhibitor at pH 6.5) was added to the column and mixed for 10 min before elution. The eluate was kept at 4 °C for further experiments.

### In vitro lipid binding assay

Protein samples were diluted to a final concentration of 9 µM with proper salt concentration (12.5 mM Na_2_HPO_4_, 125 mM NaCl). The diluted solution was incubated at room temperature (22°C) or 37°C overnight for equilibration of conformational changes. Just before imaging, phosphatidylcholine and phosphatidylserine were diluted in decane to a final concentration of 20 mM and mixed 1:1. 1.2 µl of protein solution was added to 20 µl of lipid solution in a 384 well plate (CellVis # P384-1.5H-N) followed by vibrant mixing (30-40 times) with pipettes. Samples were imaged under a confocal microscope.

### Immunoblotting

7×10^5^ cells were plated in each well of a 6 well plate, transfected using the calcium phosphate method, and incubated at the indicated temperatures. Cells were washed in PBS and lysed in RIPA buffer (50 mM Tris pH 7.5, 150 mM NaCl, 1% NP40, 0.1% SDS, 0.5% DOC, 1 mM EDTA, 2 mM sodium vanadate and protease inhibitor). 15 µL of lysate was mixed with 15µL of loading buffer (Bio-Rad #1610747) and loaded in a precast 4-15% gradient SDS-polyacrylamide gel for electrophoresis (mini-protean TGX precast gel, Bio-Rad, # 456-1084). Protein separations were transferred onto a nitrocellulose membrane using the Trans-blot Turbo RTA transfer kit (Bio-rad, #170-4270) according to manufacturer’s protocol. Membranes were blocked in 5% milk in Tris buffer saline with 0.5% Tween-20 (TBS-T) for 1 hour and incubated overnight at 4°C with primary antibodies against GFP (abcam #ab290) and tubulin (CST #3873). Each primary antibody was used at a dilution of 1:1000 in TBS-T with 3% BSA. After washing with TBS-T, membranes with incubated with secondary antibodies in TBS-T with 3% BSA for 1 hr at room temperature (IRDye^®^ 800CW Goat anti-Rabbit IgG, 1;20,000 dilution, LI-COR #926-32211; IRDye^®^ 680RD Donkey anti-Mouse IgG, 1:20,000 dilution, LI-COR, #926-68072). Membranes were then imaged on the LI-COR Odyssey scanner.

### Co-immunoprecipitation

Cells were transfected with the constructs of interest, allowed to express for 24 hrs, and subjected to the specified treatment. Subsequently, cells were washed with PBS and lysed (50 mM HEPES pH 7.4, 150 mM NaCl, 1% Triton X-100, 1 mM EDTA, 1 mM EGTA, 10% glycerol, 2 mM sodium vanadate and protease inhibitor (Sigma #P8340)). Cleared cell lysates were incubated for 2 hours with Protein A/G agarose beads (Santa Cruz, SC-2003) that were hybridized with either GFP (Thermo #GF28R) or Flag antibodies (CST #14793S). Beads were then washed 5 times with HNTG buffer (20 mM HEPES pH 7.4, 150 mM NaCl, 0.1% Triton X-100, 10% glycerol), and sample buffer was added to elute proteins. Eluates were then used for immunoblotting.

### Tissue phantom synthesis

Tissue phantoms were generated by mixing 2g of Agar powder (Fisher BP9744) in 100 µL of water and microwaving until powder was dissolved. 0.3 g Al_2_O_3_ and 0.3 mL India Ink (Pro Art PRO-4100) were then mixed in with the liquid agar and poured into a 3D printed mold designed to allow the phantom to encase an 8 well (ibidi #80826) cell culture slide. Experiments were performed by extracting the phantom from the mold, placing culture slides with cells into the solidified phantom, and subjecting the phantom/encased plate to the temperature/light exposure indicated. Illumination was performed by place the phantom on top of an optoPlate-96^57^ with the LEDs underneath the phantom programmed to be on at maximum intensity (180mW/cm^2^) and various duty cycles depending on the condition (1s On every 10s at 0mm phantom thickness and constantly On at >0mm thickness). Ambient temperature was changed by adjusting the set point of the cell culture incubator.

### Mouse maintenance

Animal experiments were performed following Protocol 807519 approved by the UPenn Institutional Animal Care and Use Committee (IACUC). NSG mice (6–8 weeks old, male) purchased from and housed by the Perelman School of Medicine Stem Cell and Xenograft Core.

### H3122 xenografts

Xenografts were performed by suspending 2×10^6^ H3122 cells expressing the indicated constructs in 100 µL of PBS+2% FBS and mixing with 100 µL of VitroGel (The Well Biosciences #VHM01). This mixture was kept in a 37°C water path while mice were prepared for injection. Mice were anesthetized using 2.5% isoflurane and 200 µL of the cell suspension was injected subcutaneously on each mouse flank. Mice were maintained under a heat lamp during injection and while recovering from anesthesia.

### Thermoelectric cooling device

The thermoelectric cooling device consists of two Peltier plates connected in series. The smaller Peltier plate (Digikey 102-4428-ND) is attached by its heating face to the cooling face of the larger Peltier plate (CNBTR TES1-4902) using thermally conductive tape (AI AIKENUO 8541602030). An electronic thermometer (Walfront MF55) is attached the cooling face of the smaller Peltier and covered with a soft thermal pad (Arctic Cooling ACTPD00004A). The thermal pad provides a soft surface when pressed against the mouse’s skin. An aluminum heat sink (Jienk JT371-374) is attached to the heating face of the larger Peltier plate to dissipate excess heat. Finally, a fan (Winsinn FAN40105V) is attached on top of the heat sink for additional heat dissipation. An Arduino microcontroller (Arduino A000053) obtains readings from the electronic thermometer and adjusts the on/off state of a transistor (Bridgold B07R49F39B) that regulates power delivery to the Peltier assembly. 3.5V is supplied to the Peltier plates when cooling is desired. The fan is constantly turned on even when no cooling is needed.

### Local cooling of mouse xenografts

Mice were anesthetized using 2.5% isoflurane, placed on a heating pad (37°C), and kept under anesthesia using a nose cone, with isoflurane percentage adjusted to maintain at least 10 breaths per 15 seconds. Local cooling was applied to the designated flank by pressing the thermoelectric cooling device to the skin with enough pressure to slightly depress the surrounding tissue.

### Luminescence imaging

Mice were injected with 200 µL of 15 mg/mL D-Luciferin (GoldBio LUCK) via intraperitoneal injection 10 minutes prior to imaging. Mice were then anesthetized with 2.5% isoflurane and luminescence was recorded using an IVIS Spectrum imaging system every ∼5 minutes until the luminescent signal was maximal. Mice were then allowed to recover from anesthesia under a heat lamp.

## Supplemental Figures

**Figure S1.**
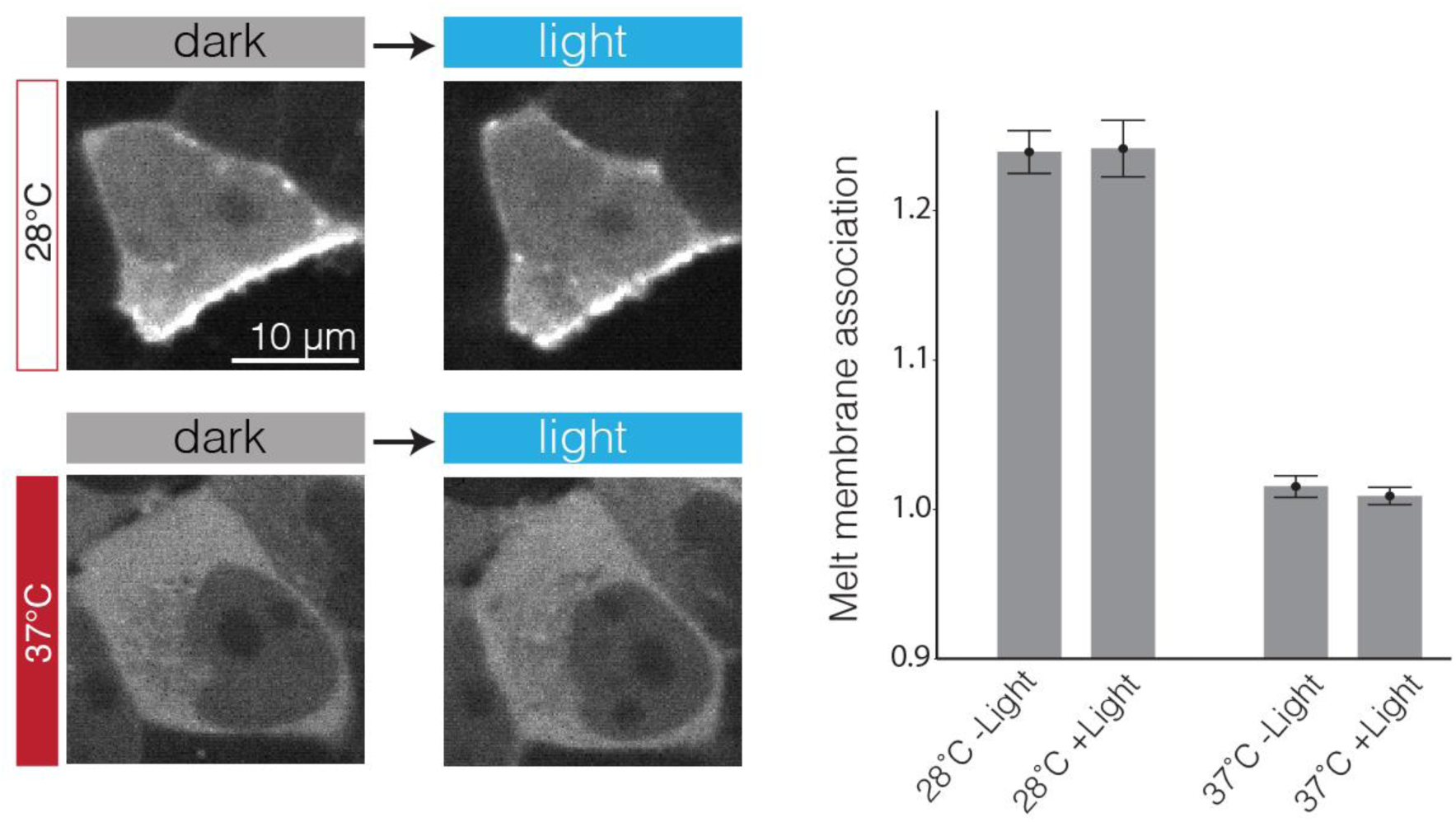
Melt is insensitive to light at high and low temperatures. To examine whether Melt was sensitive to light at either high or low temperatures, HEKs stably expressing Melt-mCh were cultured at either 28°C or 37°C for 12 hours. After 12 hours, cells were imaged, exposed to blue light for 5 min (1s 488 nm laser light every 10 s), and imaged immediately thereafter. Representative images (A) and quantification (B) and showed that light exposure did not measurably alter membrane binding under either low or high temperatures. Data in (B) represent the mean +/- SEM of ∼500 cells.

**Figure S2.**
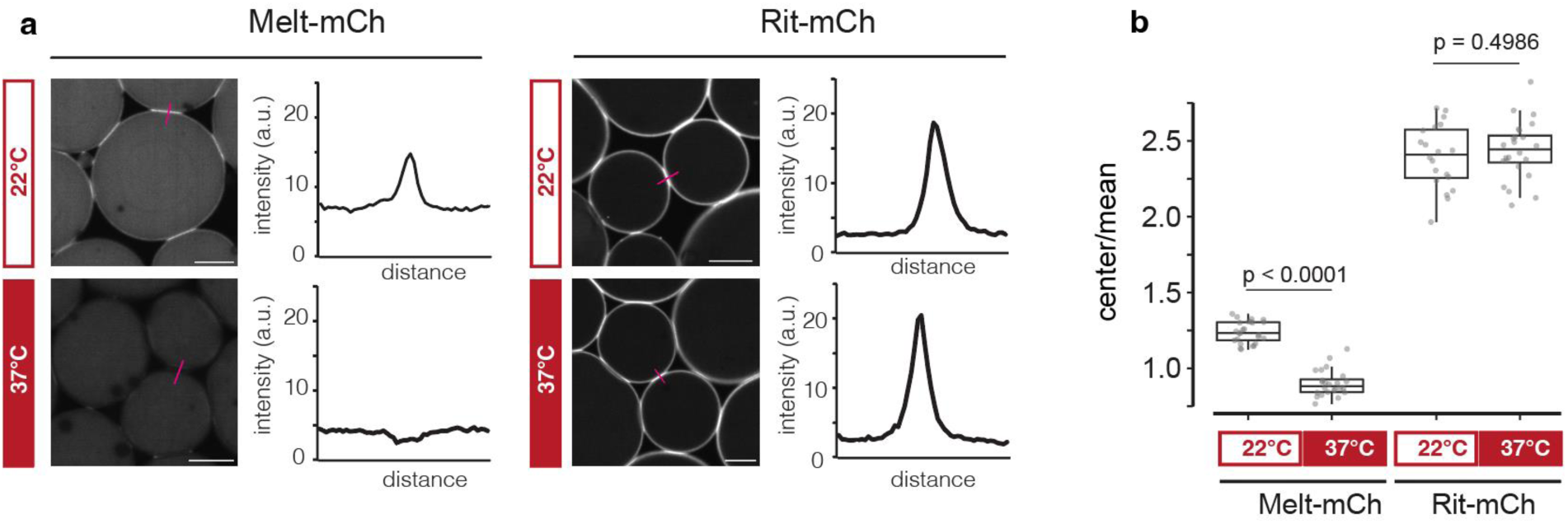
Melt shows temperature-sensitive lipid association *in vitro*. Temperature-dependent membrane binding was tested using purified protein *in vitro* and was compared to purified Rit-mCh. Rit-mCh comprises mCherry fused to the polybasic domain of Rit, which should bind to lipids in a temperature-independent manner and thus serves as a positive control of membrane binding. An aqueous solution of either Melt-mCh or Rit-mCh was incubated overnight at the designated temperatures and subsequently mixed with a lipid solution (10 mM phosphatidylserine and 10 mM phosphatidylcholine dissolved in decane). (A) Representative images and fluorescence intensity profiles over individual protocell boundaries. Scale = 5 µm. (B) Quantifications of the ratio of fluorescence on the boundary and within protocells. Each data point represents a different protocell pair (>20 pairs per condition). Significance level assessed by Students t-test. These data indicate that temperature has a direct effect on the Melt protein.

**Figure S3.**
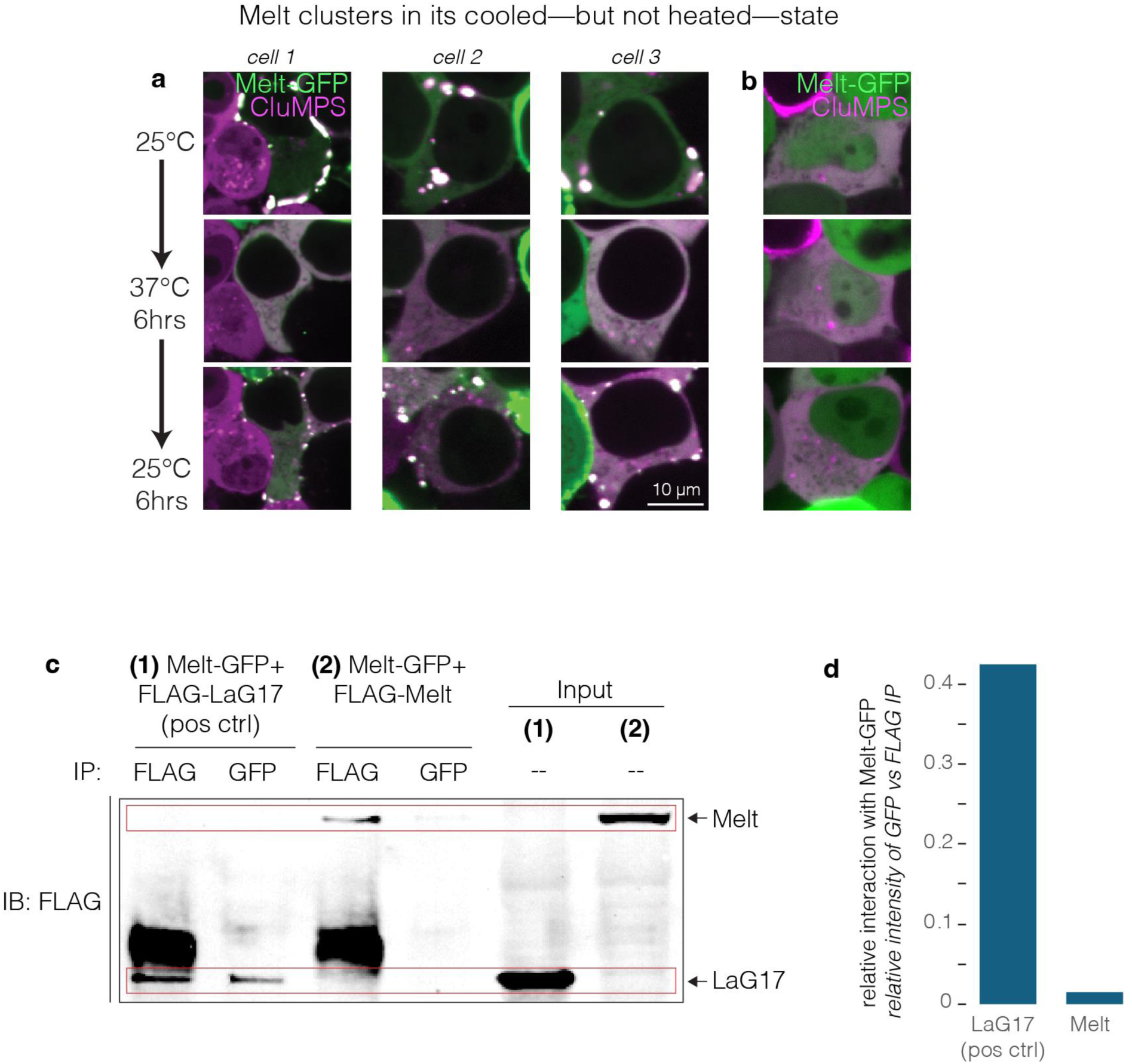
Melt multimerization is temperature sensitive. A) Examining Melt multimerization at high and low temperatures in the presence of a CluMPS reporter, which amplifies clusters of its target (in this case GFP^28^). Images show HEK 293T cells stably expressing CluMPS that were transfected with plasmid encoding Melt-GFP. CluMPS-amplified Melt clusters were present at 25°C, disappeared at 37°C, and reformed upon return to 25°C. These results demonstrate that, in addition to membrane binding, Melt clustering is also temperature sensitive. B) Negative control demonstrating that GFP alone does not cluster in the presence of CluMPS. C) Testing Melt multimerization in the heated state using co-immunoprecipitation. Previous studies suggested that BcLOV4 constitutively forms dimers/trimers *in vitro*^24^. To test if Melt dimerizes in the cytoplasm, we designed a co-IP assay where Melt was tagged with either FLAG tag or GFP, and we tested whether FLAG-tagged Melt would co-precipitate when Melt-GFP was pulled down. As a positive control, we performed co-IP on cells co-expressing Melt-GFP and FLAG-LaG17, a nanobody that binds GFP with 50 nM affinity^58^. In this positive control (1), pulldown with either FLAG or GFP allowed detection of the FLAG-LaG17 band, confirming association between the two constructs. However, in cells cotransfected with Melt-GFP and FLAG-Melt (2), pulldown of Melt-GFP revealed minimal co-precipitation with FLAG-Melt. D) Quantification of (C). Together, these data suggest that Melt is largely monomeric in its heated state in cells.

**Figure S4.**
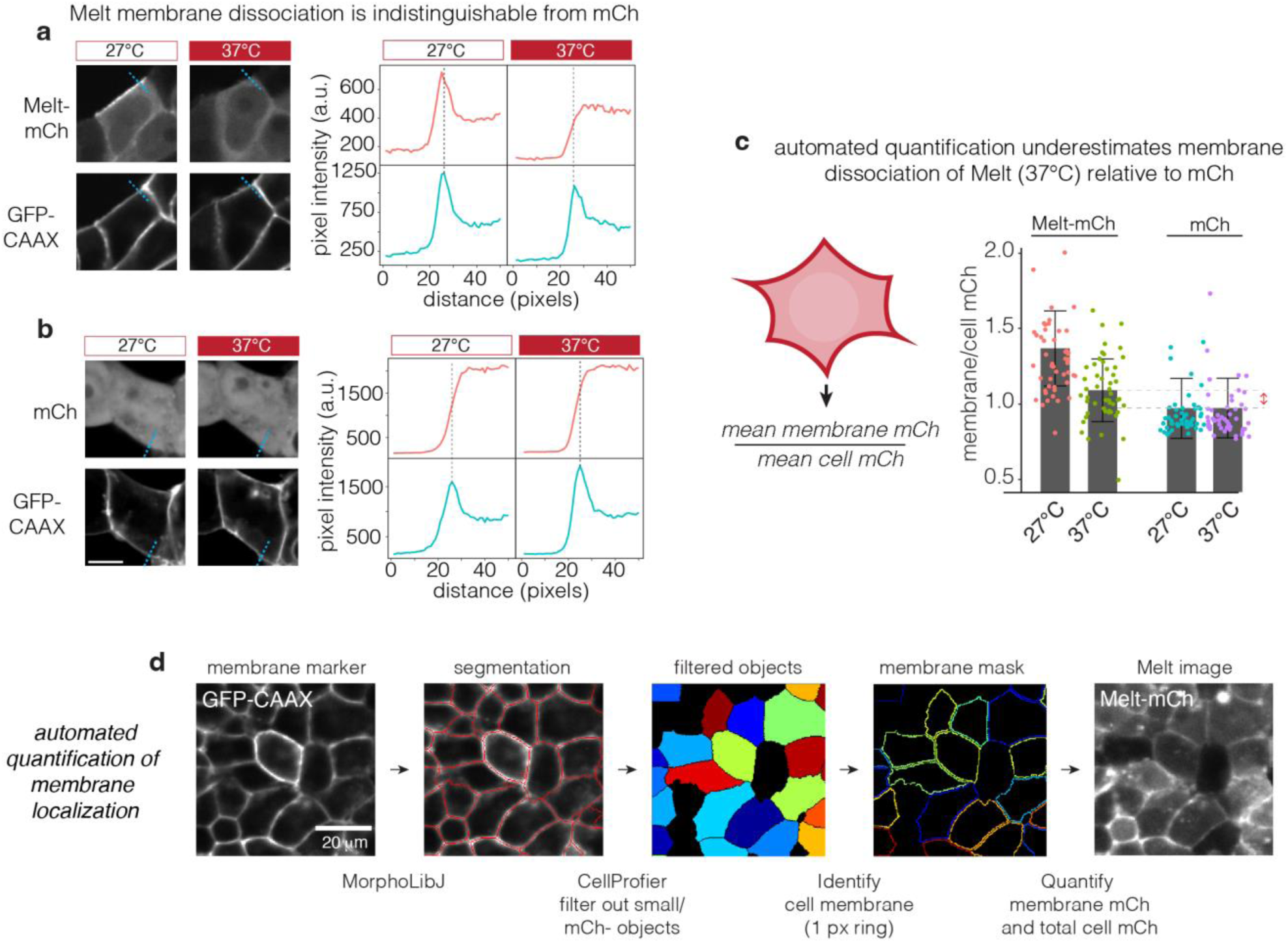
Calibrating and measuring the degree of Melt membrane association. To assess the degree of Melt membrane dissociation, we compared fluorescence profiles of Melt-mCh vs soluble mCh, which has no membrane association. A) HEK 293T cells stably expressing Melt-mCh and a GFP-CAAX membrane marker were cultured at either 27°C or 37°C for 24 hrs. Images of cells showed visible membrane binding at 27°C and no visible membrane binding at 37°C. Line profiles were taken across the membrane for both Melt-mCh and GFP-CAAX membrane marker. At the location of peak GFP-CAAX intensity, no peak is observed in the Melt-mCh channel at 37°C, indicating no residual membrane binding when Melt is temperature inactivated. B) The same quantification performed in (A) was performed with cytoplasmic soluble mCh. At both 37 and 27°C, mCh showed no peak in fluorescence across the membrane, with a profile indistinguishable from Melt in its heated state (A), thus indicating that Melt fully dissociates upon heating. C) Melt membrane association was quantified by normalizing mean membrane intensity to the mean intensity of the cell. This metric gave a slightly larger minimum value (∼1.1) than that derived from images of soluble mCh (∼1), despite their similar line profiles (A,B). This difference is likely an artifact that results from nuclear exclusion Melt, which reduces its mean cell intensity. We therefore adjusted measurements of Melt membrane association by this correction factor (1.1) throughout the manuscript so that the minimal membrane/cell fluorescence would equal 1, as observed for mCh (as justified by (A,B)) B) Illustration of automated image feature extraction. Segmentation of the GFP-CAAX membrane marker allowed high-throughput quantification of mean membrane and mean cell intensities.

**Figure S5.**
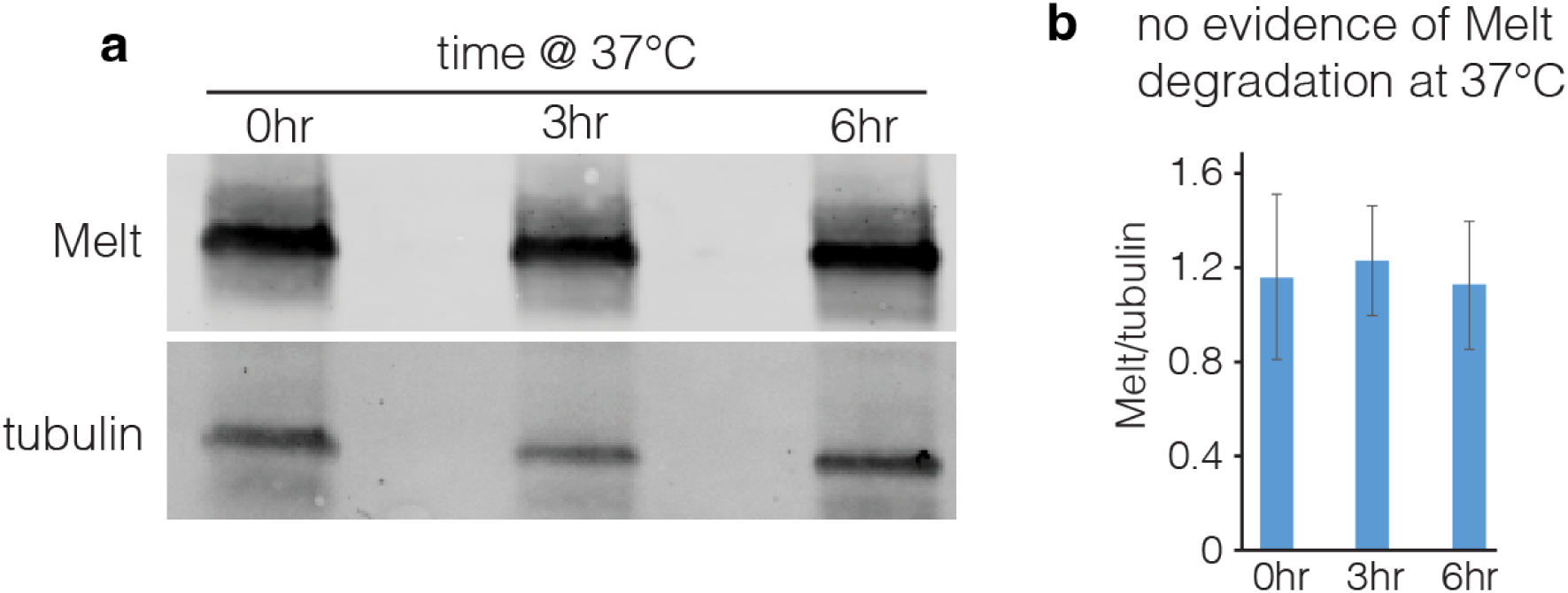
No evidence of Melt degradation in its heated state. a) Western blot from cells expressing Melt and exposed to various durations of high temperature. Cells expressing Melt-GFP were incubated at 25°C overnight, exposed to 37°C for 0, 3, or 6 hrs, and lysed. B) Densitometry from three experiments depicted in (A). No differences in protein abundance were observed. Data represent mean +/- SD of 3 experiments.

**Figure S6.**
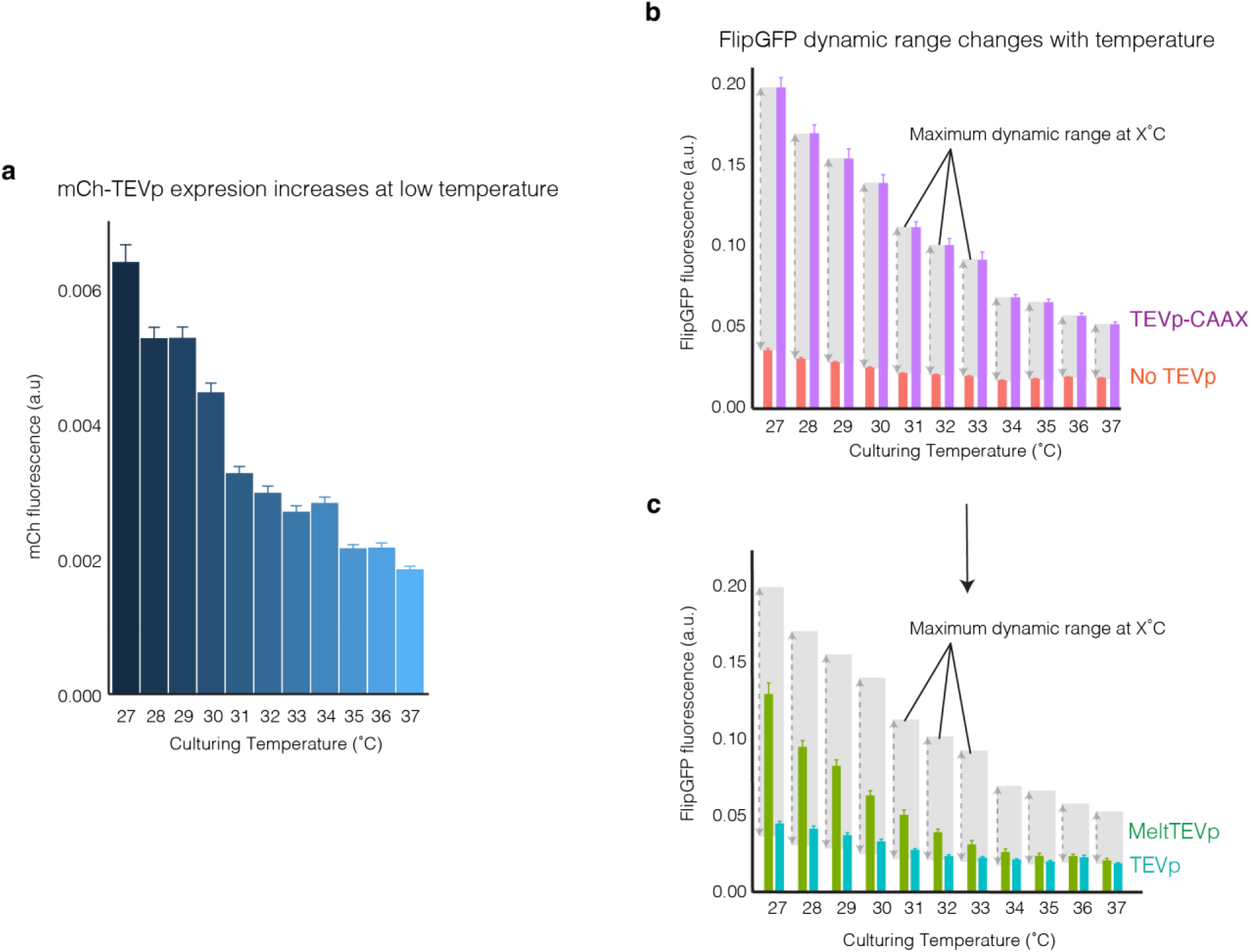
Normalization of MeltTEVp proteolysis to account for temperature-dependent changes in protein expression. A) Total protein expression is elevated at low temperatures as demonstrated by mCh-TEVp expression. Cells were cultured at the indicated temperature for 24 hours. B) To account for changes in FlipGFP signals caused by temperature-dependent expression differences, negative control (no TEVp) and positive control (constitutively membrane bound TEVp-CAAX) cells were used to establish minimal and maximal FlipGFP signals at each temperature. C) Minimal and maximal cutting ranges at each temperature were used to normalize MeltTEVp and TEVp proteolysis to the ranges established in (B) (subtracting minimum signal and dividing by maximum). This normalization was performed to account for changes in protein expression levels that resulted from increases in proteolysis at low temperatures. Each bar in all plots represents the mean +/- 1 SEM of ∼1000 cells.

**Figure S7.**
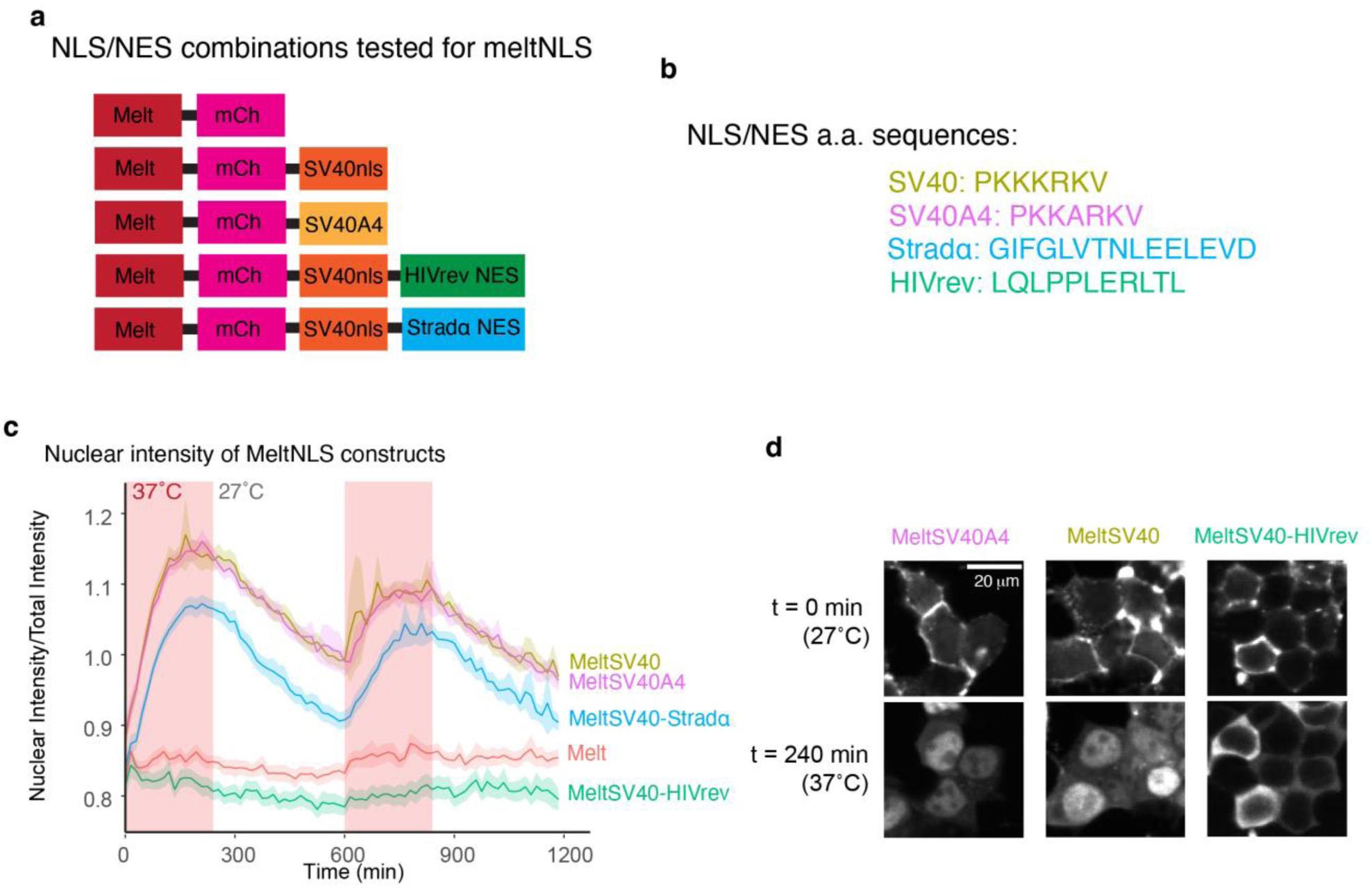
Different NLS/NES combinations achieve varying levels of nuclear shuttling. A) Diagram of all MeltNLS/NES fusions tested in order to achieve the largest dynamic range of nuclear shuttling between 27°C and 37°C. B) Amino acid sequence of NLS and NES used in MeltNLS/NES fusions. C) Quantification of nuclear Melt signal using the five constructs shown in (A) exposed to repeated cycles of heating and cooling. Traces represent the mean of ∼1000 cells +/- SEM. D) Representative images of MeltNLS/NES combinations before and after heating to 37°C and cooling to 27°C.

**Figure S8.**
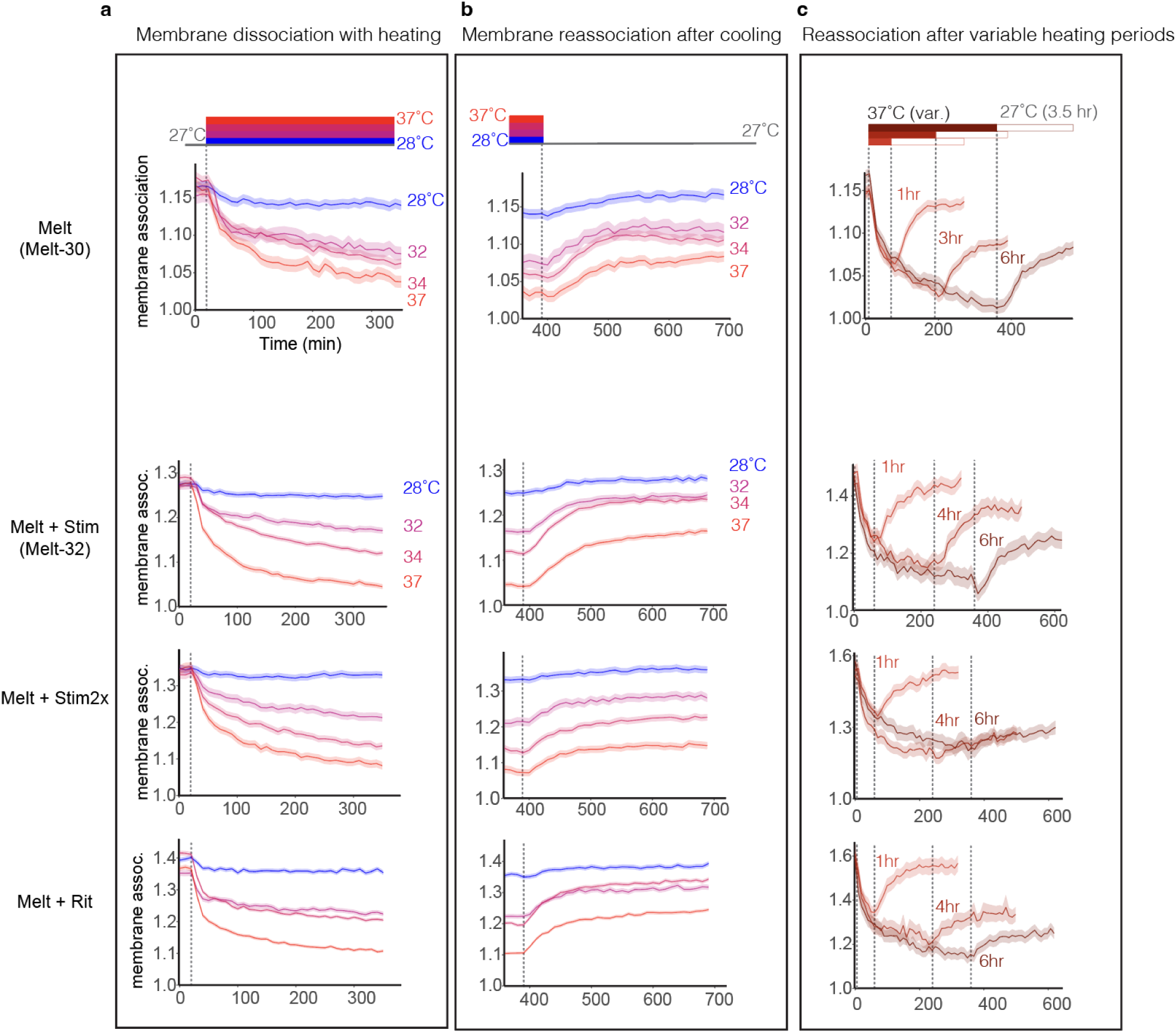
Kinetics of membrane dissociation and reassociation of Melt-PB fusions. A) Quantification of membrane dissociation at the indicated temperature after prior culture at 27°C for 24 hours. Dashed lines indicate the time at which the temperature was raised to the indicated temperature. B) Quantification of membrane recruitment of the indicated construct cultured at 27°C after previous culture at the indicated temperature for the preceding 6 hours. Traces represent the kinetics of membrane reassociation and are continuations of traces found in (A). Dashed lines indicate the time at which the temperature was lowered from the indicated temperature. C) Quantification of membrane recruitment of the indicated construct during culture at 37°C following culture at 27°C for 24 hours. Dashed lines indicate the time at which cells were returned to 27°C to identify the effect of different periods of heating on membrane reassociation kinetics. All traces represent the mean of ∼1000 cells +/- SEM.

**Figure S9.**
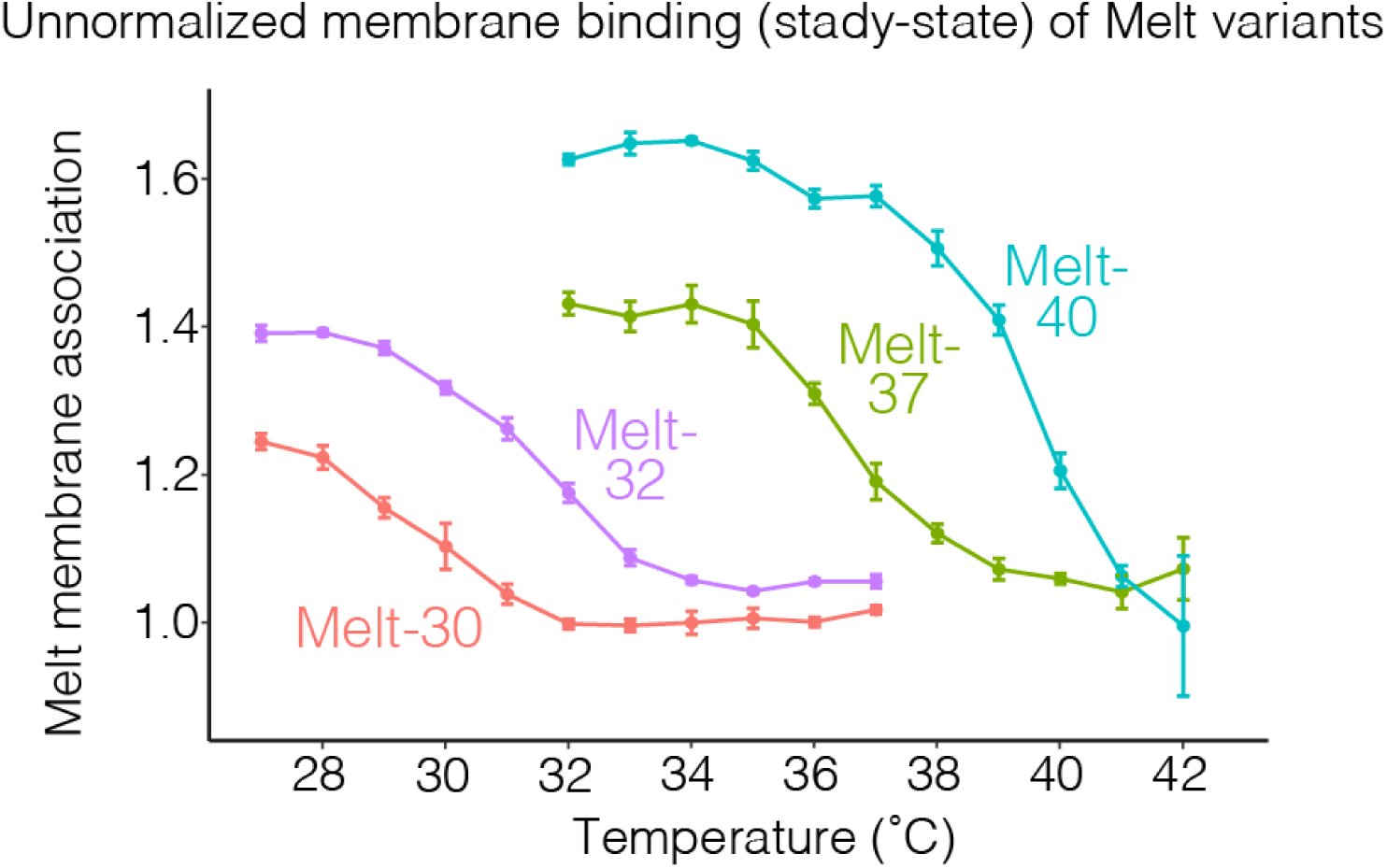
Relative membrane binding of Melt variants. Unnormalized plots of data shown in **Figure 4H**, showing relative membrane binding strength of Melt-30/32/37/40 at the indicated temperatures. Each point represents the average of three wells +/- SD with ∼500 cells quantified in each well.

**Figure S10.**
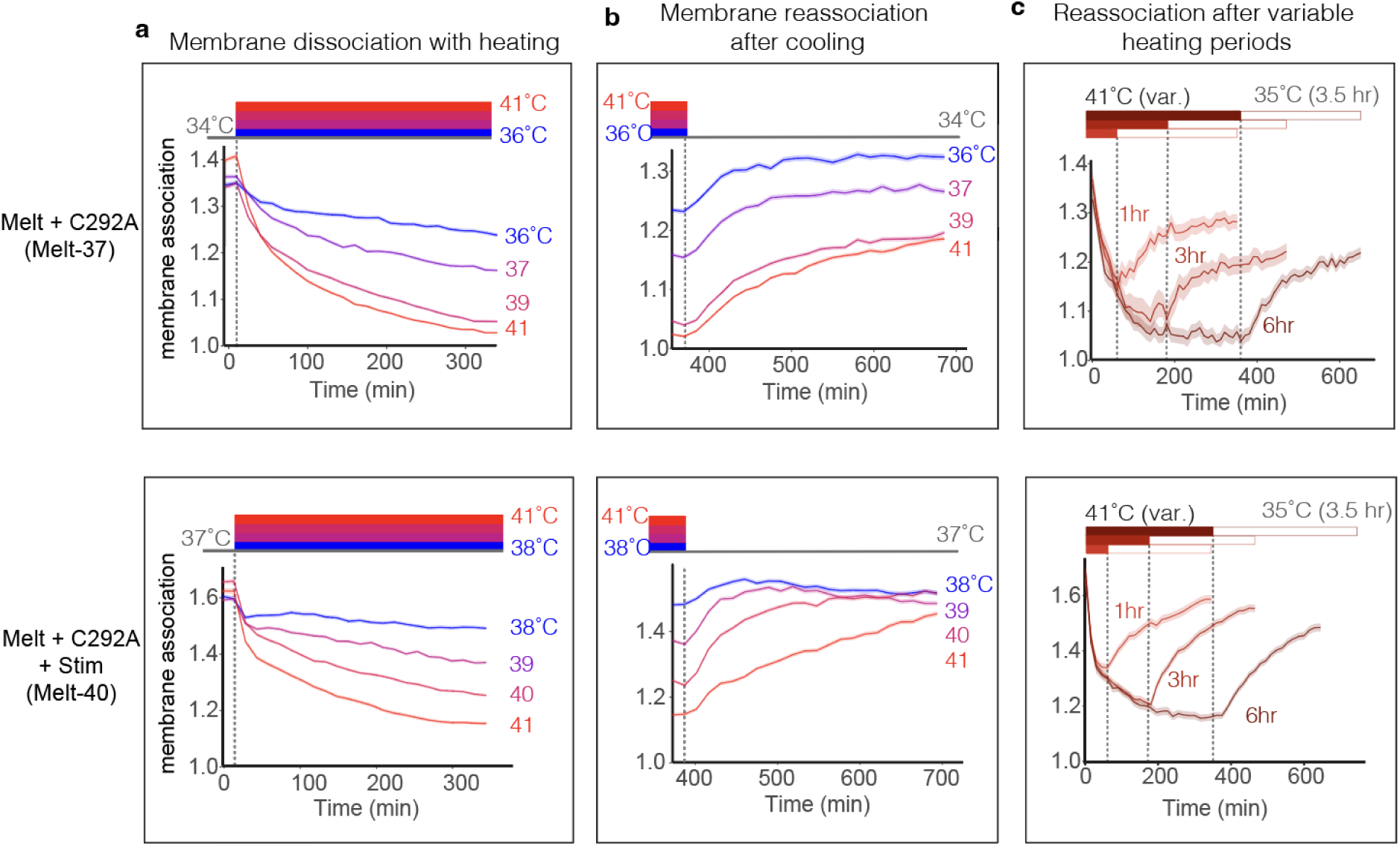
Kinetics of membrane dissociation and reassociation of Melt variants. A) Quantification of membrane recruitment of the indicated construct cultured at the indicated temperatures. Traces represent the kinetics of membrane dissociation after prior culture at either 34°C (C292A) or 37°C (C292A+Stim) for 24 hours. Dashed lines indicate the time at which the temperature was raised to the indicated temperature. B) Quantification of membrane recruitment of the indicated construct cultured at 34°C (C292A) or 37°C (C292A+Stim) after prior culture at the indicated temperature for the preceding 6 hours. Traces represent the kinetics of membrane reassociation and are continuations of traces found in (A). Dashed lines indicate the time at which the temperature was lowered from the indicated temperature. C) Quantification of membrane recruitment of the indicated construct during culture at 41°C after prior culture at 35°C for 24 hours. Dashed lines indicate the time at which cells were returned to 35°C to identify the effect of different periods of heating on membrane reassociation kinetics. All traces represent the mean +/- SEM of ∼1000 cells.

**Figure S11.**
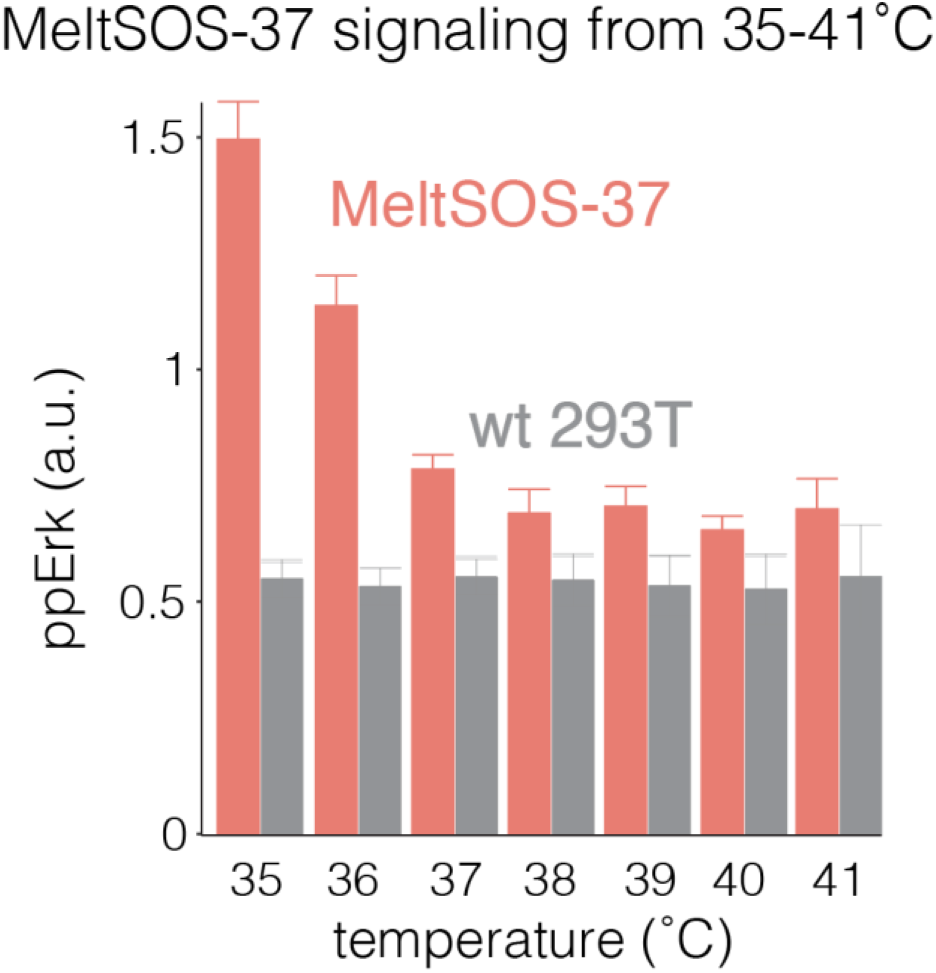
Thermal activation of MeltSOS-37. MeltSOS-37 achieves signaling activation at temperatures < 37°C. Plot showing quantification of pathway activation (single-cell immunofluorescence for ppErk) in cells expressing MeltSOS-37 exposed to the indicated temperatures for 75 min. Data points represent the mean of 2 wells +/- SD with ∼1000 cells quantified per well.

**Figure S12.**
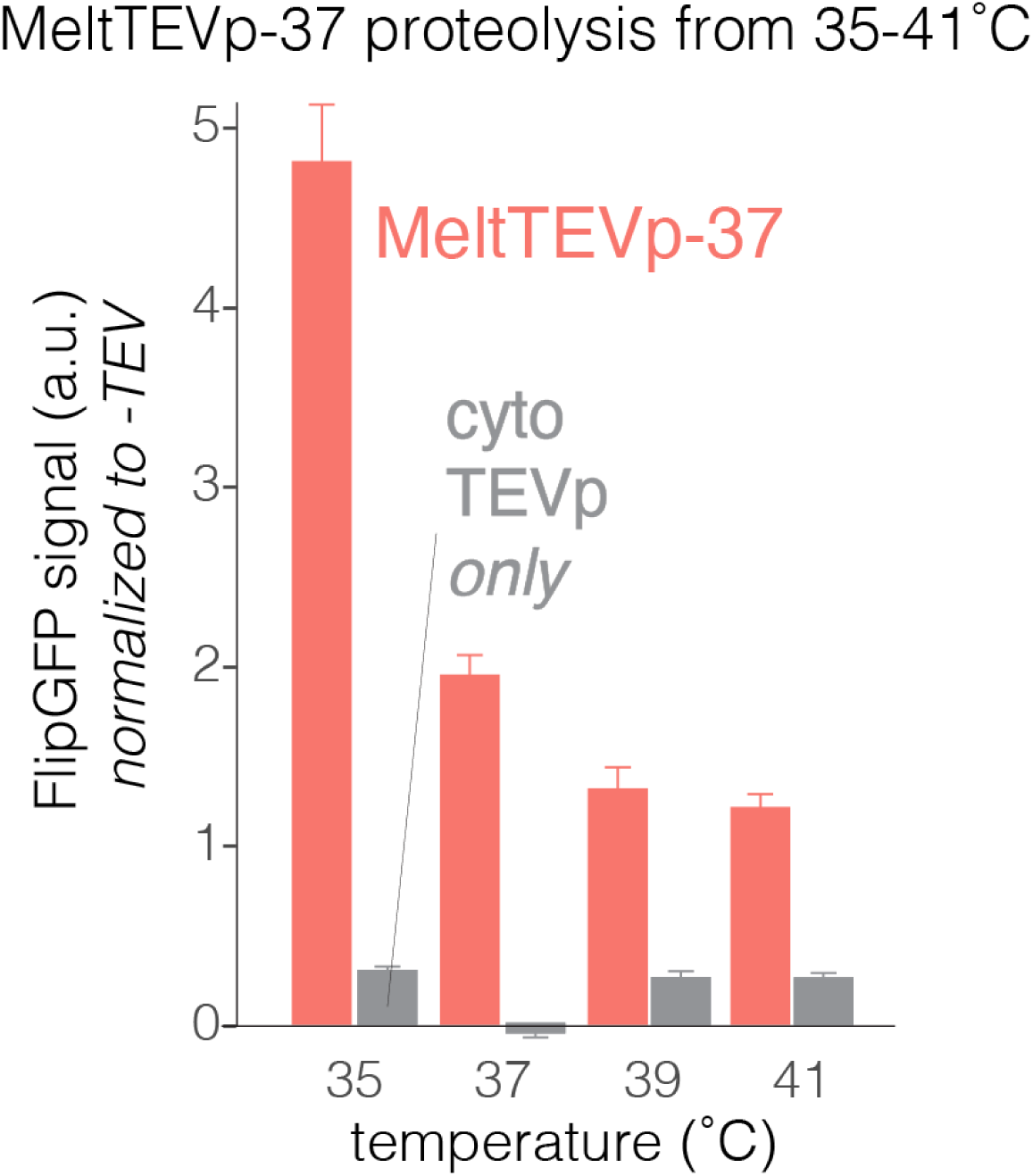
Thermal activation of MeltTEVp-37. MeltTEVp-37 achieves proteolysis at temperatures <37°C. Plot showing FlipGFP fluorescence in cells expressing MeltTEVp exposed to the indicated temperatures. Data points represent the mean ∼1000 cells +/- SEM. See **Methods** for FlipGFP quantification workflow.

**Figure S13.**
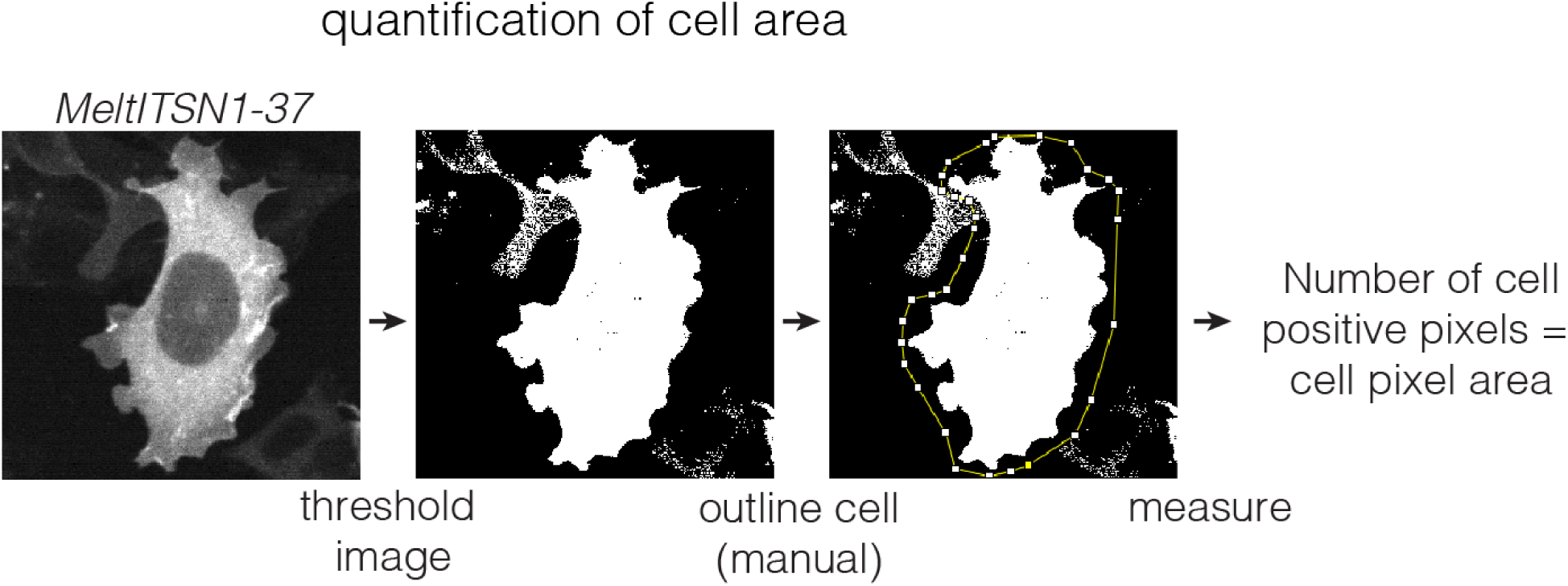
Quantification of cell area to assess effects of meltITSN1-37. A cell expressing MeltITSN1-37 was imaged and subsequently thresholded in ImageJ such that pixels within the cell were set to 1 and background pixels were set to 0. A region of interest containing the cell of interest was drawn by hand. Summing the total number of positive pixels in the cell region was used as a metric of total cell area.

**Figure S14.**
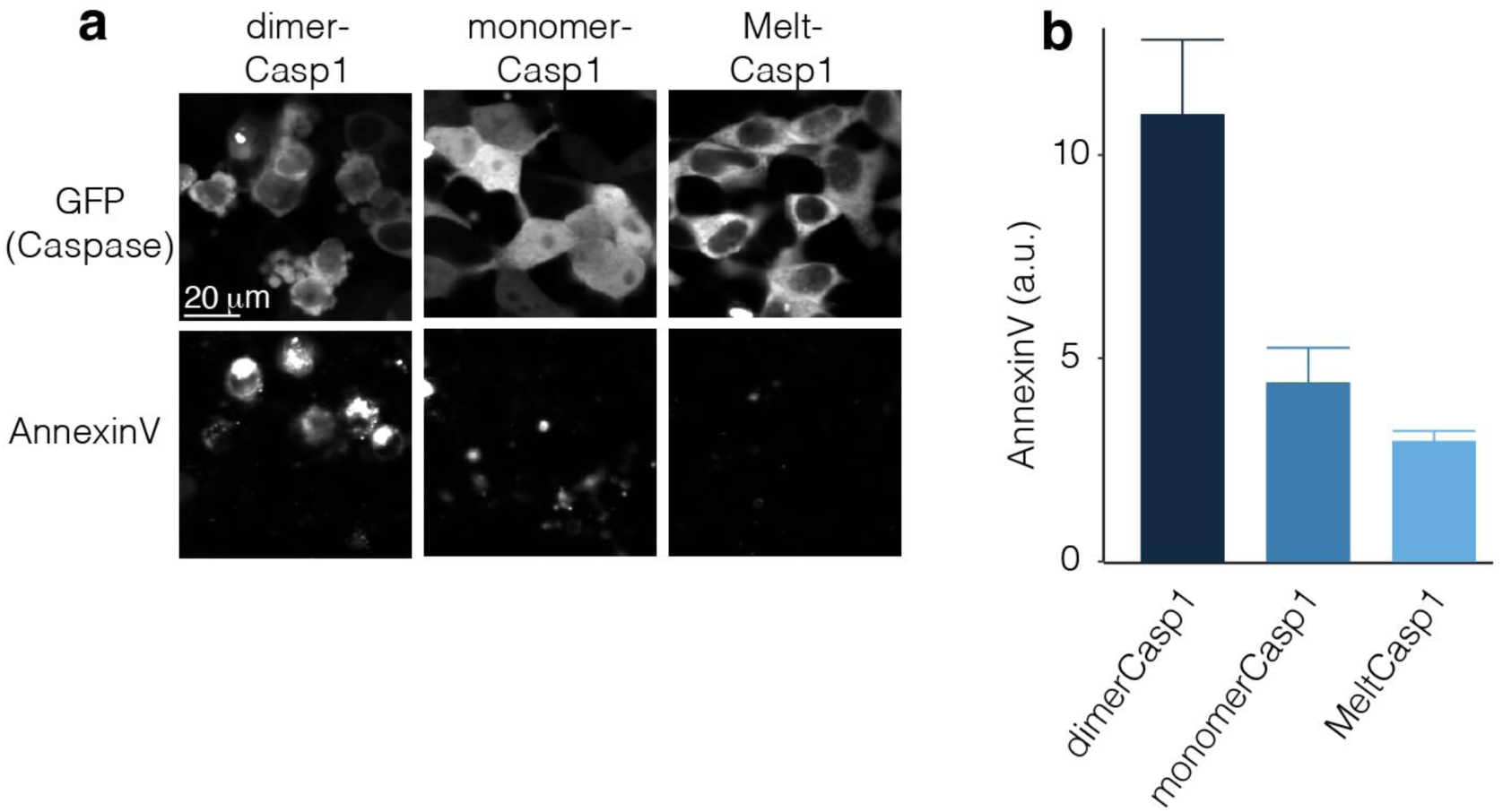
MeltCasp1 demonstrates low background cell death. A) Representative images of HEK 293T cells transiently transfected with either GCN4p1-caspase1 (dimer), caspase1 (monomer), or MeltCaspase1-30 fused to GFP. The dimer was sufficient to drive noticeable cell death relative to the monomeric caspase, as measured both by cell morphology and Annexin V staining. Cell death in MeltCasp1-30-expressing cells maintained at 37°C was comparable to the monomer-Casp1, further indicating a lack of Melt self-association in its heated state. B) Quantification of Annexin V from the experiment in (A). Annexin levels in MeltCasp-30 cells was comparable to monomeric caspase-1 and substantially lower than the dimeric construct. Data represent the mean +/- 1 SEM of three wells.

**Figure S15.**
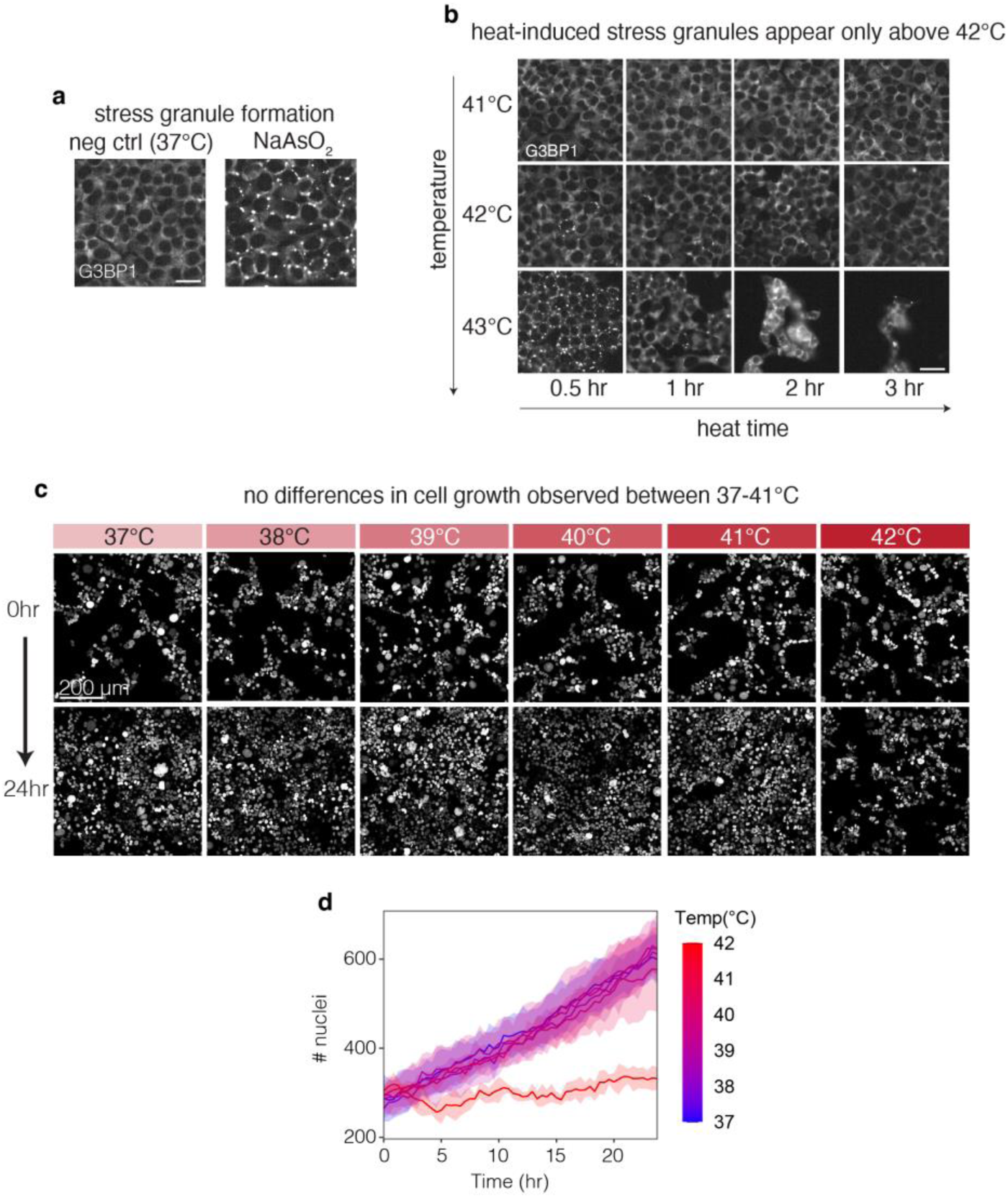
Lack of thermal stress observed below 42°C. To examine whether the temperature changes required for Melt-37/40 activation would also apply thermal stress to mammalian cells, we measured stress granule (SG) formation as well as changes in proliferation in response to thermal stimuli used throughout the manuscript. A) SGs were visualized by immunofluorescence for G3BP1. No SGs were seen in HEK 293T cells in normal growth conditions, while bright SG puncta were seen in cells treated with 100 µM sodium arsenite for 3 hours prior to fixation (positive control). B) SGs were visualized in HEK 293Ts that were exposed to various durations and intensities of heating. No SGs were observed in cells heated to ≤ 41°C, and only a few cells showed SGs when heated to 42°C. By contrast, heating to 43°C induced SGs in nearly all cells within 30 min, followed by detachment of cells at later time points. C) To examine integration of potential heat stress over longer time periods, we measured growth and proliferation of HEK 293T cells grown at temperatures between 37-42°C over 24 hours. Images show cell nuclei (H2B-iRFP) at T = 0 and T = 24 hrs. D) Quantification of (C) shows that growth appeared unperturbed between 37-41°C, with a dramatic reduction in proliferation at 42°C. Traces represent the mean +/- SD of 4 imaging fields.

**Figure S16.**
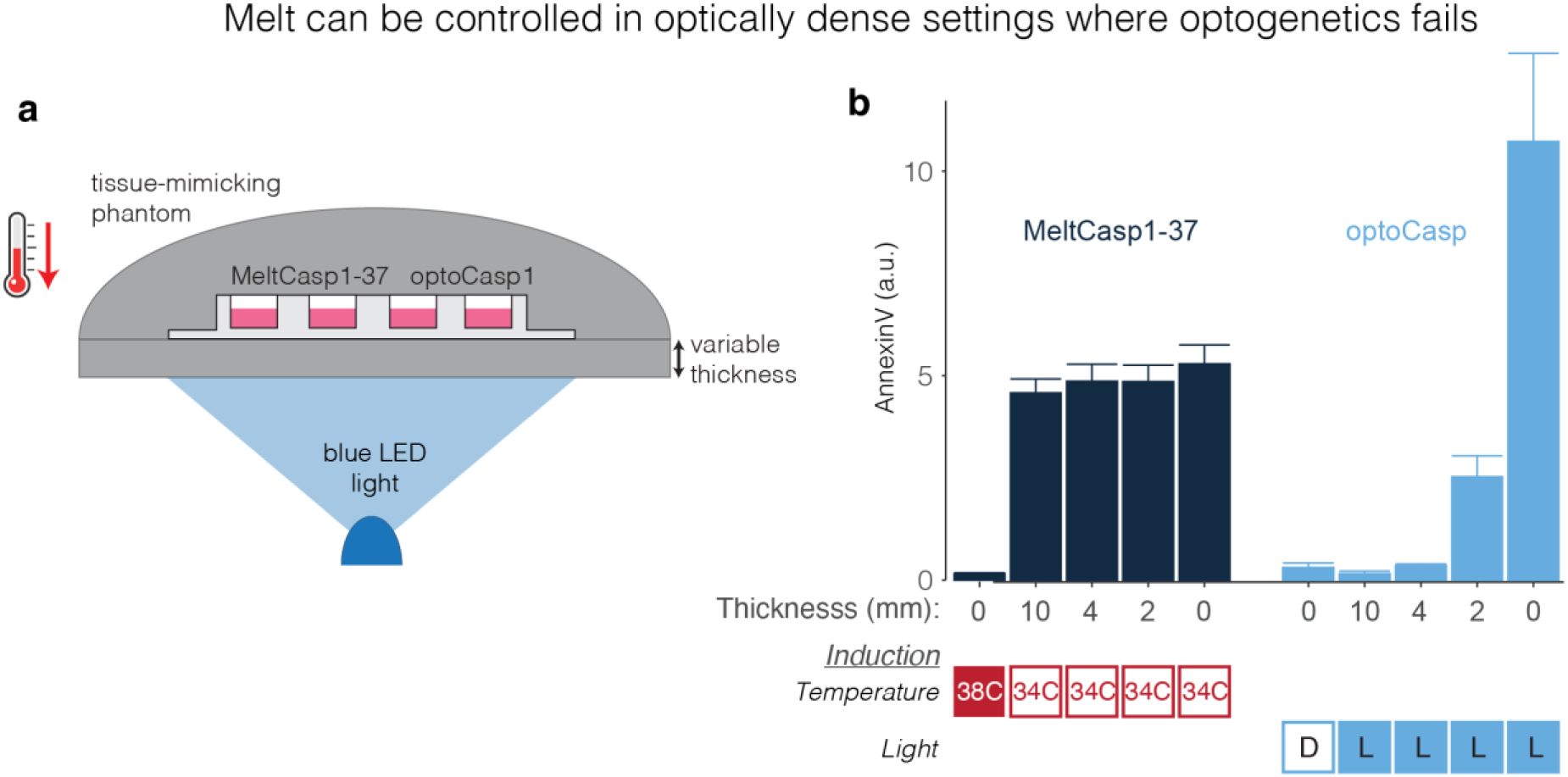
Melt functions in opaque settings. A) Schematic of experimental setup. Tissue phantoms were generated (see **Methods**) that mimic the light and temperature absorption properties of human tissue. Phantoms were used to enclose cultures of HEK cells expressing either MeltCasp1-37 or optoCasp1. The bottom of the phantom was adjusted to a thickness ranging from 0 to 10 mm. B) MeltCasp1-37 could be actuated independent of phantom thickness by adjusting the ambient temperature. At 0 mm thickness, optoCasp1 showed robust cell death when exposed to blue light. However, at a thickness of 2 mm, optoCasp1 induction was significantly attenuated and completely abolished at 4mm. Data points represent the mean +/- SEM of three wells at 8 hours post-induction (light or heat). The optoPlate-96^57^ was used for blue light exposure with cells receiving 1s of light every 10s at 0 mm thickness and constant light at > 0 mm thickness. See **Methods** for further details.

**Figure S17.**
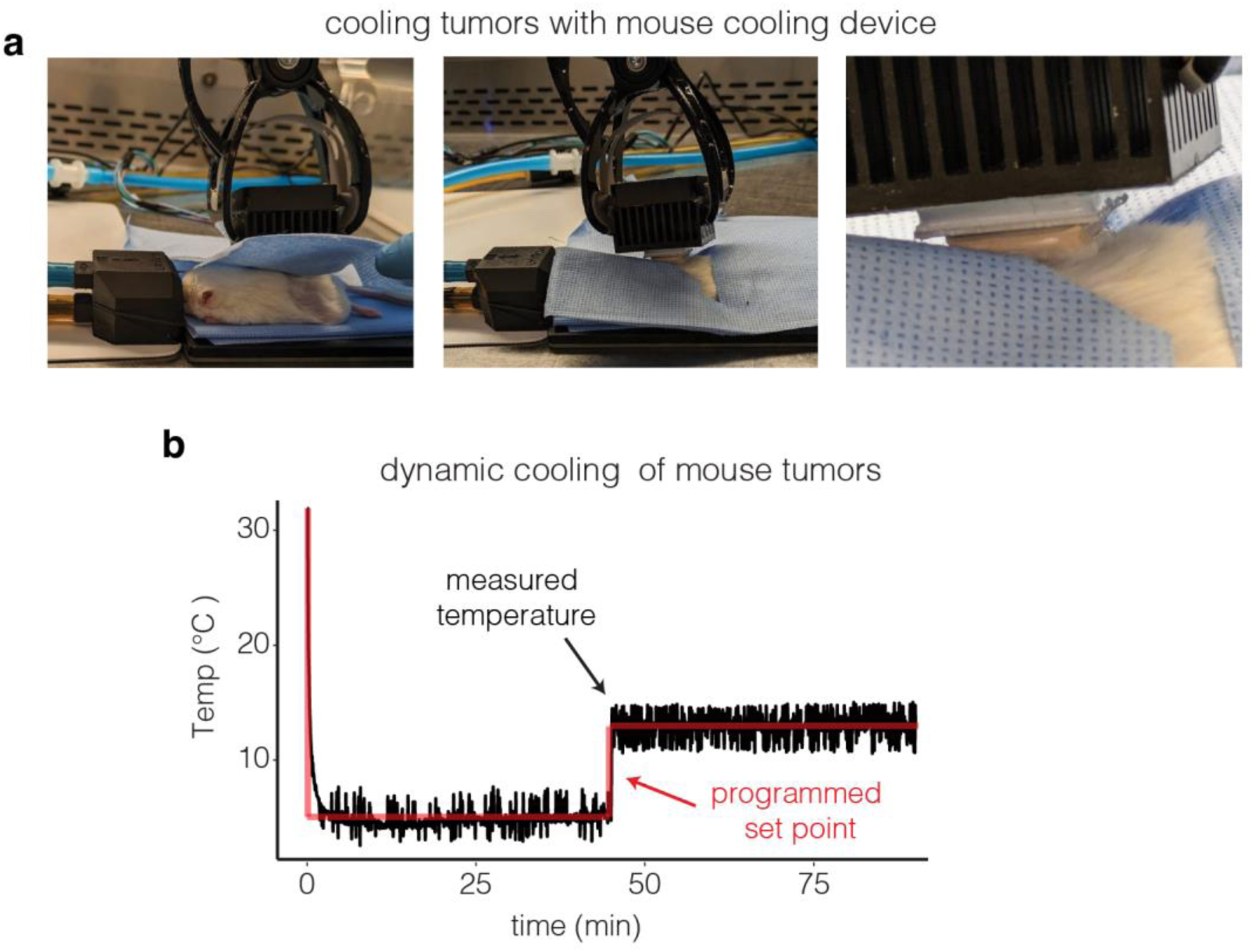
Topical application of cooling to mice. A) Images of mouse undergoing localized cooling. Mice were kept under constant anesthetization on top of a heating pad and underneath a surgical blanket. A hole in the surgical blanket allows contact between the cooling device and the area of the mouse targeted for cooling. B) Temperature readings from the thermometer at the contact between the device and the mouse were recorded and plotted. The device was able to rapidly lower the interface temperature to both desired setpoints (5°C and 15°C), dynamically transitioning between them during the experiment.

## Supplementary Movie Captions

**Supplementary Movie 1. Reversible membrane binding of Melt using temperature changes.** HEK 293T cells stably expressing Melt were exposed to 1 hour of heating followed by 4 hours of cooling (37° and 27°C respectively) in order to capture dynamic changes in membrane binding at each temperature. Time is hh:mm. Scale bar = 40 µm.

**Supplementary Movie 2. Temperature-controlled nucleocytoplasmic shuttling of MeltNLS/NES.** HEK 293T cells transiently expressing MeltNLS/NES were exposed to repeated rounds of 37° and 27°C to observe dynamic changes in nuclear shuttling. Time is hh:mm. Scale bar = 15 µm.

**Supplementary Movie 3. Thermal control of Erk activity in mammalian temperature ranges using MeltEGFR-37.** HEK 293T cells stably expressing MeltEGFR-37 were exposed to repeated rounds of 37° and 40°C. Video shows the ErkKTR reporter, which indicates Erk activation through changes in the ratio of cytoplasmic to nuclear fluorescence. Nuclear enrichment of the reporter upon heating indicates reduction of Ras-Erk signaling, while nuclear depletion upon cooling indicates pathway activation. Stills from this movie were used to generate the images found in **Figure 4K**. Time is hh:mm. Scale bar = 10 µm.

**Supplementary Movie 4. Temperature-controlled nucleocytoplasmic shuttling of MeltNLS/NES-40 in mammalian temperature ranges.** HEK 293T cells transiently expressing MeltNLS/NES-40 were exposed to repeated rounds of 41° and 37°C in order to capture dynamic changes in nuclear shuttling. Time is hh:mm. Scale bar = 20 µm.

**Supplementary Movie 5. Reversible changes in cell size through thermal control of MeltITSN1-37.** Cells expressing MeltITSN1-37 were cultured at 41°C for 24 hours prior to imaging. Upon lowering the temperature to 37°C, cells showed rapid expansion in size, which could be toggled over multiple rounds of heating and cooling. Time is hh:mm. Scale bar = 20 µm.

**Supplementary Movie 6. Temperature-inducible cell death using MeltCasp1-37.** HEK 293T cells transiently expressing MeltCasp1-37 were exposed to either maintained 38°C or cooled to at 34°C. Cells cooled to 34°C showed morphological changes associated with apoptosis, increased Annexin V staining, and detachment from the plate. Time is hh:mm. Scale bar = 40 µm. MeltCasp1-37 is shown in green while Annexin V-647 is shown in magenta.

## References and Notes

1. Ntziachristos, V. Going deeper than microscopy: the optical imaging frontier in biology. Nat. Methods 7, 603–614 (2010).

2. Ash, C., Dubec, M., Donne, K. & Bashford, T. Effect of wavelength and beam width on penetration in light-tissue interaction using computational methods. Lasers Med. Sci. 32, 1909–1918 (2017).

3. Piraner, D. I. et al. Going Deeper: Biomolecular Tools for Acoustic and Magnetic Imaging and Control of Cellular Function. Biochemistry 56, 5202–5209 (2017).

4. Miller, I. C. et al. Enhanced intratumoural activity of CAR T cells engineered to produce immunomodulators under photothermal control. Nat Biomed Eng 5, 1348–1359 (2021).

5. Ermakova, Y. G. et al. Thermogenetic control of Ca2+ levels in cells and tissues. bioRxiv 2023.03.22.533774 (2023) doi:10.1101/2023.03.22.533774.

6. Corbett, D. C. et al. Thermofluidic heat exchangers for actuation of transcription in artificial tissues. Sci Adv 6, (2020).

7. Walton, M., Roestenburg, M., Hallwright, S. & Sutherland, J. C. Effects of ice packs on tissue temperatures at various depths before and after quadriceps hematoma: Studies using sheep. J. Orthop. Sports Phys. Ther. 8, 294–300 (1986).

8. ter Haar, >gail & Coussios, C. High intensity focused ultrasound: Physical principles and devices. Int. J. Hyperthermia 23, 89–104 (2007).

9. Horowitz, N. H. Biochemical Genetics of Neurospora. in Advances in Genetics (ed. Demerec, M.) vol. 3 33–71 (Academic Press, 1950).

10. Talavera, A. & Basilico, C. Temperature sensitive mutants of BHK cells affected in cell cycle progression. J. Cell. Physiol. 92, 425–436 (1977).

11. Varadarajan, R., Nagarajaram, H. A. & Ramakrishnan, C. A procedure for the prediction of temperature-sensitive mutants of a globular protein based solely on the amino acid sequence. Proc. Natl. Acad. Sci. U. S. A. 93, 13908–13913 (1996).

12. Hurme, R., Berndt, K. D., Normark, S. J. & Rhen, M. A proteinaceous gene regulatory thermometer in Salmonella. Cell 90, 55–64 (1997).

13. Piraner, D. I., Wu, Y. & Shapiro, M. G. Modular Thermal Control of Protein Dimerization. ACS Synth. Biol. 8, 2256–2262 (2019).

14. Guo, Y., Liu, S., Jing, D., Liu, N. & Luo, X. The construction of elastin-like polypeptides and their applications in drug delivery system and tissue repair. J. Nanobiotechnology 21, (2023).

15. Despanie, J., Dhandhukia, J. P., Hamm-Alvarez, S. F. & MacKay, J. A. Elastin-like polypeptides: Therapeutic applications for an emerging class of nanomedicines. J. Control. Release 240, 93–108 (2016).

16. Li, Z., Tyrpak, D. R., Park, M., Okamoto, C. T. & MacKay, J. A. A new temperature-dependent strategy to modulate the epidermal growth factor receptor. Biomaterials 183, 319–330 (2018).

17. Abedi, M. H., Lee, J., Piraner, D. I. & Shapiro, M. G. Thermal Control of Engineered T-cells. ACS Synth. Biol. 9, 1941–1950 (2020).

18. Wu, Y. et al. Control of the activity of CAR-T cells within tumours via focused ultrasound. Nat Biomed Eng 5, 1336–1347 (2021).

19. Morimoto, R. I. Cells in stress: transcriptional activation of heat shock genes. Science 259, 1409–1410 (1993).

20. Feder, M. E. & Hofmann, G. E. HEAT-SHOCK PROTEINS, MOLECULAR CHAPERONES, AND THE STRESS RESPONSE: Evolutionary and Ecological Physiology. (2003) doi:10.1146/annurev.physiol.61.1.243.

21. Akerfelt, M., Morimoto, R. I. & Sistonen, L. Heat shock factors: integrators of cell stress, development and lifespan. Nat. Rev. Mol. Cell Biol. 11, 545–555 (2010).

22. Benman, W. et al. Temperature-responsive optogenetic probes of cell signaling. Nat. Chem. Biol. 18, 152–160 (2022).

23. Benman, W., Iyengar, P., Mumford, T., Huang, Z. & Bugaj, L. J. Multiplexed dynamic control of temperature to probe and observe mammalian cells. bioRxiv (2024) doi:10.1101/2024.02.18.580877.

24. Glantz, S. T. et al. Directly light-regulated binding of RGS-LOV photoreceptors to anionic membrane phospholipids. Proc. Natl. Acad. Sci. U. S. A. 115, E7720–E7727 (2018).

25. Pal, A. A., Benman, W., Mumford, T. R., Chow, B. Y. & Bugaj, L. J. Optogenetic clustering and membrane translocation of the BcLOV4 photoreceptor. bioRxiv 2022.12.12.520131 (2022) doi:10.1101/2022.12.12.520131.

26. Harper, S. M., Neil, L. C. & Gardner, K. H. Structural basis of a phototropin light switch. Science 301, 1541–1544 (2003).

27. Huang, Z., Benman, W., Dong, L. & Bugaj, L. J. Rapid optogenetic clustering in the cytoplasm with BcLOVclust. J. Mol. Biol. 168452 (2024).

28. Mumford, T. R. et al. Simple visualization of submicroscopic protein clusters with a phase-separation-based fluorescent reporter. Cell Syst. 15, 166–179.e7 (2024).

29. Grecco, H. E., Schmick, M. & Bastiaens, P. I. H. Signaling from the living plasma membrane. Cell 144, 897–909 (2011).

30. Toettcher, J. E., Weiner, O. D. & Lim, W. A. Using optogenetics to interrogate the dynamic control of signal transmission by the Ras/Erk module. Cell 155, 1422–1434 (2013).

31. Citri, A. & Yarden, Y. EGF-ERBB signalling: towards the systems level. Nat. Rev. Mol. Cell Biol. 7, 505–516 (2006).

32. Liang, S. I. et al. Phosphorylated EGFR Dimers Are Not Sufficient to Activate Ras. Cell Rep. 22, 2593–2600 (2018).

33. Chung, H. K. et al. A compact synthetic pathway rewires cancer signaling to therapeutic effector release. Science 364, (2019).

34. Gao, X. J., Chong, L. S., Kim, M. S. & Elowitz, M. B. Programmable protein circuits in living cells. Science 361, 1252–1258 (2018).

35. Sanchez, M. I. & Ting, A. Y. Directed evolution improves the catalytic efficiency of TEV protease. Nat. Methods 17, 167–174 (2020).

36. Zhang, Q. et al. Designing a Green Fluorogenic Protease Reporter by Flipping a Beta Strand of GFP for Imaging Apoptosis in Animals. J. Am. Chem. Soc. 141, 4526–4530 (2019).

37. Collas, P. & Aleström, P. Nuclear localization signal of SV40 T antigen directs import of plasmid DNA into sea urchin male pronuclei in vitro. Mol. Reprod. Dev. 45, 431–438 (1996).

38. Dorfman, J. & Macara, I. G. STRADalpha regulates LKB1 localization by blocking access to importin-alpha, and by association with Crm1 and exportin-7. Mol. Biol. Cell 19, 1614–1626 (2008).

39. Heo, W. D. et al. PI(3,4,5)P3 and PI(4,5)P2 lipids target proteins with polybasic clusters to the plasma membrane. Science 314, 1458–1461 (2006).

40. He, L. et al. Optical control of membrane tethering and interorganellar communication at nanoscales. Chem. Sci. 8, 5275–5281 (2017).

41. Regot, S., Hughey, J. J., Bajar, B. T., Carrasco, S. & Covert, M. W. High-Sensitivity Measurements of Multiple Kinase Activities in Live Single Cells. Cell 157, 1724–1734 (2014).

42. Levskaya, A., Weiner, O. D., Lim, W. A. & Voigt, C. A. Spatiotemporal control of cell signalling using a light-switchable protein interaction. Nature 461, 997–1001 (2009).

43. Berlew, E. E. et al. Designing Single-Component Optogenetic Membrane Recruitment Systems: The Rho-Family GTPase Signaling Toolbox. ACS Synth. Biol. 11, 515–521 (2022).

44. Hannanta-Anan, P., Glantz, S. T. & Chow, B. Y. Optically inducible membrane recruitment and signaling systems. Curr. Opin. Struct. Biol. 57, 84–92 (2019).

45. Nobes, C. D. & Hall, A. Rho, rac, and cdc42 GTPases regulate the assembly of multimolecular focal complexes associated with actin stress fibers, lamellipodia, and filopodia. Cell 81, 53–62 (1995).

46. Shkarina, K. et al. Optogenetic activators of apoptosis, necroptosis, and pyroptosis. J. Cell Biol. 221, (2022).

47. Gallouzi, I. E. et al. HuR binding to cytoplasmic mRNA is perturbed by heat shock. Proc. Natl. Acad. Sci. U. S. A. 97, 3073–3078 (2000).

48. Kedersha, N. L., Gupta, M., Li, W., Miller, I. & Anderson, P. RNA-binding proteins Tia-1 and tiar link the phosphorylation of eIF-2α to the assembly of mammalian stress granules. J. Cell Biol. 147, 1431–1442 (1999).

49. Ntombela, L., Adeleye, B. & Chetty, N. Low-cost fabrication of optical tissue phantoms for use in biomedical imaging. Heliyon 6, e03602 (2020).

50. Berlew, E. E. et al. Single-component optogenetic tools for inducible RhoA GTPase signaling. Advanced Biology 2100810 (2021).

51. Berlew, E. E., Kuznetsov, I. A., Yamada, K., Bugaj, L. J. & Chow, B. Y. Optogenetic Rac1 engineered from membrane lipid-binding RGS-LOV for inducible lamellipodia formation. Photochem. Photobiol. Sci. (2020) doi:10.1039/c9pp00434c.

52. Qiao, J., Peng, H. & Dong, B. Development and Application of an Optogenetic Manipulation System to Suppress Actomyosin Activity in Ciona Epidermis. Int. J. Mol. Sci. 24, (2023).

53. Wang, W. et al. A light-and calcium-gated transcription factor for imaging and manipulating activated neurons. Nat. Biotechnol. 35, 864–871 (2017).

54. Tidyr. https://tidyr.tidyverse.org/.

55. Wickham, H. ggplot2. WIREs Computational Statistics 3, 180–185 (2011).

56. Legland, D., Arganda-Carreras, I. & Andrey, P. MorphoLibJ: integrated library and plugins for mathematical morphology with ImageJ. Bioinformatics 32, 3532–3534 (2016).

57. Bugaj, L. J. & Lim, W. A. High-throughput multicolor optogenetics in microwell plates. Nat. Protoc. 14, 2205–2228 (2019).

58. Fridy, P. C. et al. A robust pipeline for rapid production of versatile nanobody repertoires. Nat. Methods 11, 1253–1260 (2014).

